# Construction of a bioluminescence-based assay for bitter taste receptors (TAS2Rs)

**DOI:** 10.1101/2021.11.07.467592

**Authors:** Shi Min Tan, Wei-Guang Seetoh

## Abstract

In humans, a family of 25 bitter taste receptors (TAS2Rs) mediates bitter taste perception. A common approach to characterize bitter causative agents involves expressing TAS2Rs and the appropriate signal transducers in heterologous cell systems, and monitoring changes in the intracellular free calcium levels upon ligand exposure using a fluorescence-based modality, which typically suffers from a low signal window, and is susceptible to interference by autofluorescence, therefore prohibiting its application to screening of plant or food extracts, which are likely to contain autofluorescent compounds. Here, we report the development and validation of a bioluminescence-based intracellular calcium release assay for TAS2Rs that has a better assay performance than a fluorescence-based assay. Furthermore, the bioluminescence-based assay enables the evaluation of TAS2R agonists within an autofluorescent matrix, highlighting its potential utility in the assessment of the bitterness-inducing properties of plant or food fractions by the food industry. Additionally, improvement to the bioluminescence-based assay for some TAS2Rs was achieved by altering their N-terminal signal sequences, leading to signal window enhancement. Altogether, the bioluminescence-based TAS2R assay can be used to perform functional studies of TAS2Rs, evaluate TAS2R-modulating properties of autofluorescent samples, and facilitate the discovery of compounds that can function as promising bitter taste modulators.

## Introduction

The gustatory system in humans enables the perception of five basic taste sensations (sweet, umami, sour, salty, and bitter) and allows us to assess the nutritive, hedonic, and safety qualities of food and beverages. A class of 25 bitter taste receptors (TAS2Rs) mediates bitter taste perception in humans. TAS2Rs are expressed in Type II taste receptor cells and belong to the G-protein coupled receptor (GPCR) superfamily.^1^ However, TAS2Rs have also been identified in extra-oral tissues, such as the heart, lungs, and intestines, which implicate their involvement in various physiological and pathological processes and position some TAS2Rs as potential therapeutic targets.^2^ TAS2Rs possess highly diverse coding sequences, sharing 16-84% sequence homology in the transmembrane segments.^3^ Furthermore, genetic variation across the human TAS2R repertoire adds complexity to bitter taste perception and shapes individual differences in bitter taste sensitivity and food preferences.^4^ A bitter tastant can serve as an agonist of several TAS2Rs, and a single TAS2R can bind several structurally diverse bitter ligands.^5^ Binding of an agonist to a TAS2R induces conformational changes in the receptor, which are transduced by heterotrimeric G-proteins and amplified by second messenger systems to effect cellular responses.

Several techniques can be used to monitor TAS2R activation, but cell-based functional assays that measure events distal to receptor activation (e.g., changes in second messenger levels) are commonly used because amplification of the signaling pathway is involved, which boosts the signal-to-background ratio of the assay. Significant TAS2R studies typically employ an *in vitro* assay system that involves a TAS2R engineered for cell surface expression, and a chimeric Gα protein. Changes in TAS2R activation prompt transient fluctuations in intracellular calcium levels, which can be measured using a fluorescent calcium indicator (e.g., Fluo-4, Calcium 6) whose emission intensity varies in direct proportion to the free calcium concentration. Even though fluorometric analyses have been widely employed in TAS2R assays to identify the functional effects of natural products and synthetic ligands, it is challenging to evaluate the potential bitter modulatory effects of complex biological samples. Extracts and fractionates of plants and food usually contain endogenous, or processing-derived, autofluorescent compounds that will interfere with the measurement of fluorescence signals, and raise the false positive and false negative hit rates in agonist and antagonist screening campaigns respectively, with subsequent processing of false hits resulting in higher resource consumption.^8^

Alternatively, ligand-induced calcium mobilization may be assessed using calcium-sensitive photoproteins, such as aequorin and clytin, which emit bioluminescence upon binding to free calcium ions.^9, 10^ The active photoprotein complex is reconstituted from the association of coelenterazine, a luminophore, with the apophotoprotein in the presence of molecular oxygen. Binding of calcium ions to the active photoprotein eventuates in oxidative decarboxylation of coelenterazine to coelenteramide, with simultaneous release of carbon dioxide and flash emission of blue light.^11^ Photoproteins are compelling alternatives to fluorescent probes because bioluminescent responses are not affected by autofluorescence. Importantly, the luminescent intensity is directly commensurate with the free calcium concentration, and photoprotein-based assays have been validated to generate data comparable to those acquired from assays that employ fluorescent indicators.^12, 13^ Achieving a high signal-to-noise ratio in a bioluminescence-based assay is possible because cells display negligible background autoluminescence. In addition, photoproteins are amenable to recombinant expression, and genetic manipulation for targeting to specific cellular compartments such as the mitochondria to enable them to be more sensitive to changes in the free calcium levels.^14^ While functional photoprotein-based assays have been developed for various GPCRs, there is however scarce implementation of the photoprotein technology in the field of taste receptors. Hitherto, there has only been one report on the development of a bioluminescence-based assay for the human sweet taste receptor (TAS1R2/TAS1R3) that utilized mitochondrial-targeted (mt)-clytin II as the photoprotein.^12^

Here, we describe the construction of a bioluminescence-based TAS2R assay that is amenable to the screening of TAS2R agonists in an autofluorescent matrix, and has a better assay performance compared to existing fluorescence-based TAS2R assays. Further improvement to the bioluminescence-based assay for some TAS2Rs was accomplished by altering their N-terminal signal sequences, which in turn enhanced the functional expression of TAS2Rs at the plasma membrane. This approach can be readily applied to high-throughput screening assays to facilitate the discovery of compounds that can function as promising bitter taste modulators, or potential therapeutic agents.

## Methods and Materials

### Chemicals

All compounds were purchased from commercial sources. Denatonium benzoate (>98.0%), epigallocatechin gallate (>98.0%), N-acetylthiourea (>98.0%), phenylthiocarbamide (>98.0%), phenyl β-D-glucopyranoside (>98.0%), picrotoxinin (>94.0%), sinigrin (>98.0%), and stevioside (>85.0%) were purchased from Tokyo Chemical Industry. Aloin (≥95%), andrographolide (≥98%), aristolochic acid (≥98%), cromolyn sodium (≥98%), flufenamic acid (≥98%), oxyphenonium bromide (≥98%), pirenzepine ( 98%), propylthiouracil (≥ 98%), and salicin (≥98%) were purchased from Cayman Chemical. Arbutin (≥98%), brucine sulfate heptahydrate (≥98%), (98%), chloroquine diphosphate (98.5-101.0%), dimethyl thioformamide ( 98%), helicin (99%), magnesium chloride hexahydrate (≥98%), strychnine (≥98%), and zinc sulfate heptahydrate (99%) were purchased from Sigma-Aldrich. Amarogentin (98.96%) was purchased from MedChemExpress. Calcium chloride dihydrate (99.0-103.0%) was purchased from Kanto Kagaku. Restriction enzymes and Cre recombinase were purchased from New England BioLabs.

### cDNA sequences

Canonical *Homo sapiens* TAS2R protein sequences as denoted in UniProt were employed, except for TAS2R31 and TAS2R38 for which the WMVI and PAV polymorphisms were used, respectively. The protein sequences of the TAS2Rs used in this study are indicated in Supplementary Table 1.

### Construction of multigene expression vectors

Vectors purchased from the MultiBacMam kit (Geneva Biotech) were used to generate multigene vectors. The genes for mt-clytin II and *Homo sapiens* Gα16-gust44 chimera were codon-optimized for expression in human cell lines and Gα16-gust44 and mt-clytin II were cloned into CAG-promoter based donor vectors bearing the spectinomycin and kanamycin resistance gene, respectively. Each TAS2R gene was modified by replacing the initiator ATG codon with a sequence coding for the appropriate signal sequence (SST_3_ or M_3_), and cloned into the CMV-promoter based acceptor vector bearing the gentamycin resistance gene. The generation of a multigene expression vector was achieved by fusing donor and acceptor vectors using an *in vitro* Cre-*loxP* reaction by following the manufacturer’s protocol as described in the MultiBacMam kit (Geneva Biotech). g of each plasmid was mixed in a final reaction volume of 20 μL Cre recombinase, and incubated for 1 hour at 37 °C to increase the probability of forming a 1:1:1 integrated gene product. The Cre-*loxP* reaction mixture was used to transform *E. coli* DH5α by heat shock, plated onto g mL^-1^ gentamycin, 50 μg mL^-1^ kanamycin, and 100 μg mL spectinomycin after a 4 h recovery period, and allowed overnight growth at 37 °C. Several colonies were selected for overnight growth at 37 °C in LB media supplemented with the abovementioned antibiotics to purify their plasmids for restriction enzyme analysis using *Bst*XI. Based on their distinctive digestion patterns, type 1 and type 2 plasmids can be differentiated, as well as to confirm single copy integration of each plasmid in the multigene product. Two assay controls were used in this study: cells transfected with i) mt-clytin II only, and ii) both Gα16-gust44 and mt-clytin II. Only multigene vectors with single copy integration of each constituent vector were used in this study.

### Bioluminescence-based intracellular calcium release assay

293AD (Cell Biolabs, Inc) and AD-293 cells (Agilent) were maintained at 37 °C in a humidified atmosphere of 8% CO_2_ and cultured in high-glucose Dulbecco’s modified Eagle’s medium (Gibco) supplemented with 10% (v/v) heat-inactivated fetal bovine serum (Biowest) and 1% (v/v) penicillin-streptomycin (Gibco). Cells were seeded at a density of 8,400 cells per well in white 384-well tissue culture plates (Greiner) and grown overnight. After 24 h, cells were transiently transfected with 25 ng of the triple gene construct per well using ViaFect (Promega), employing a transfection agent to plasmid ratio of 3:1 μL:μg. After 24 h post-transfection, the media in each plate was replaced with 40 μL of 1× Hank’s Balanced Salt Solution (HBSS) containing 20 mM M coelenterazine h (AAT Bioquest). The assay plate was incubated at 27 °C in the dark for 4 h. Compound stock solutions were prepared by dissolving compounds in either 1× HBSS containing 20 mM HEPES pH 7.4 (buffer A), or dimethyl sulfoxide (DMSO). Ligands dissolved in DMSO were further diluted to the appropriate concentration in buffer A or LB broth, and the final DMSO concentration in the assay did not exceed 0.5% (v/v). The luminescence mode of FLIPR^TETRA^ (Molecular Devices) was employed using a gain of 28,000, and an exposure time of 1.5-3.0 s. From the source plate, 12.5 μL of test compound was dispensed into the assay plate. Each interval of luminescence signal measurement spanned 3.1 s, starting with a baseline reading of 3 intervals, and 40 intervals of signal record after ligand addition. The area under the curve (AUC) was computed using ScreenWorks (Molecular Devices), with the third interval serving as the baseline for background subtraction. Data reported were derived from at least two independent experiments performed in at least duplicates and non-linear regression analysis was performed using the 3-parameter logistic regression model in Prism 7 (GraphPad). Dose-response curves were scaled to zero baseline.

### Fluorescence-based intracellular calcium release assay

Experiments were performed as described above for the bioluminescence-based intracellular calcium release assay, except cells were seeded in black 384-well tissue culture plates (Greiner). After 24 h post-transfection, the media in each plate was replaced with 40 μL of 1× Hank’s Balanced Salt Solution (HBSS) containing 20 mM HEPES pH 7.4 and 4× dilution of Fluo4 (Thermo Fisher). The assay plate was incubated at 37 °C, 8% CO_2_ in the dark for 2 h. The fluorescence mode of FLIPR^TETRA^ was employed using an excitation of 470-495 nm and emission of 515-μL of test compound was dispensed into the assay plate. Each interval of fluorescence emission signal measurement spanned 0.63 s, starting with a baseline reading of 10 intervals, and 300 intervals of signal record after ligand addition. The difference between the maximum and minimum RFU was used to calculate dose-response curves. Data reported were derived from at least two independent experiments performed in technical quadruplicates and non-linear regression analysis was performed using the 3-parameter logistic regression model in Prism 7 (GraphPad).

### Cell surface expression HiBiT assay

A library of TAS2R20/38/50 constructs, containing N-terminal signal sequences from various Class A GPCRs followed by a HiBiT tag flanked by EFGGGSGGSSSGG and GGSGGGGSGGSSSGGVD linkers, was generated by cloning into a modified pACEMam1 vector purchased from the MultiBacMam kit (Geneva Biotech) (see Supplementary Tables 2-3, and Supplementary Figure 1 for more details). Transfection of AD-293 was performed in white 384-well tissue culture plates (Greiner) with 18 ng of each construct using ViaFect (Promega), employing a transfection agent to plasmid ratio of 3:1 μL:μg. After 24 h, the plates were equilibrated to room temperature and the culture media was replaced by the Nano-Glo HiBiT extracellular reagent (Promega), which was prepared according to the manufacturer’s protocol. After 10 minutes of incubation with the reagent at room temperature, luminescence signals were measured in a microplate reader using an integration time of 1 sec (EnSpire). After having identified that the M_3_ signal sequence generated the greatest extent of cell surface expression, we constructed the M_3_-TAS2R20/38/50 vector as described in the “Construction of multigene expression vectors” section, the sequence of which is similar to that of SST_3_-TAS2Rs in Supplementary Table 1, except that SST_3_ signal sequence is replaced by the M_3_ amino acid sequence (Supplementary Table 2). Subsequently, M_3_-TAS2R20/38/50 was used to generate the multigene vectors for functional studies using *in vitro* Cre-*loxP* reaction as described above.

## Results

### Vector construction and selection of the optimal vector type for subsequent studies

Functional TAS2R assays require the proper trafficking of receptors to the plasma membrane, and that TAS2Rs efficiently engage a second messenger system subsequent to ligand binding to produce detectable signals. Our approach to the establishment of a bioluminescence-based intracellular calcium release assay for TAS2Rs in a heterologous mammalian cell system requires three genes to be expressed: i) full-length TAS2R bearing an N-terminal rat somatostatin receptor type 3 (SST_3_) signal sequence to promote receptor translocation to the plasma membrane, ii) Gα16-gust44 chimera for functional coupling of activated TAS2R with phospholipase C activity to eventuate in the release of calcium ions from the endoplasmic reticulum (ER), and iii) mt-clytin II, a mitochondrial-targeted, calcium-binding photoprotein, which reports on TAS2R stimulation by producing bioluminescence upon binding to released calcium ions.

Ensuring heterologous expression of the three genes in a uniform manner is crucial to the development and validation of a biochemical assay for TAS2Rs. The co-transfection of three expression vectors, each encoding a different gene, is expected to be associated with significant drawbacks such as inconsistent co-transfection efficiency, and an inability to achieve a homogenous population of cells that are transfected with all three genes. Co-transfection would result in a heterogeneous population of singly and multiply (e.g. doubly and triply) transfected cells. Furthermore, triply transfected cells may not contain each of the three different plasmids. These disadvantages could potentially result in poor assay reproducibility.

Here, we relied on the approach of generating a single multigene expression plasmid encoding all three genes, with each gene placed under the control of its constitutive promoter, to ensure that transfected cells express the complete ensemble of genes that are necessary for the bioluminescence-based assay to function.^15^ Individual expression vectors encoding each of the three genes were constructed, and assembled into a singular plasmid by *in vitro* Cre-*loxP*-mediated recombination. When there is single copy integration of each vector, the recombination reaction yields two possibilities of a 17 kb plasmid construct, each with a specific pattern of gene assembly, which we assign as type 1 and type 2 (Figure 1a). To avoid multiple copy integration that may sometimes arise, it is crucial to use approximately equal amounts of each plasmid during the Cre-*loxP* reaction to increase the probability of forming the 1:1:1 integrated gene product, and verify single copy integration of the multigene vector product using restriction enzyme digestion analysis (see Methods).

**Figure 1.**
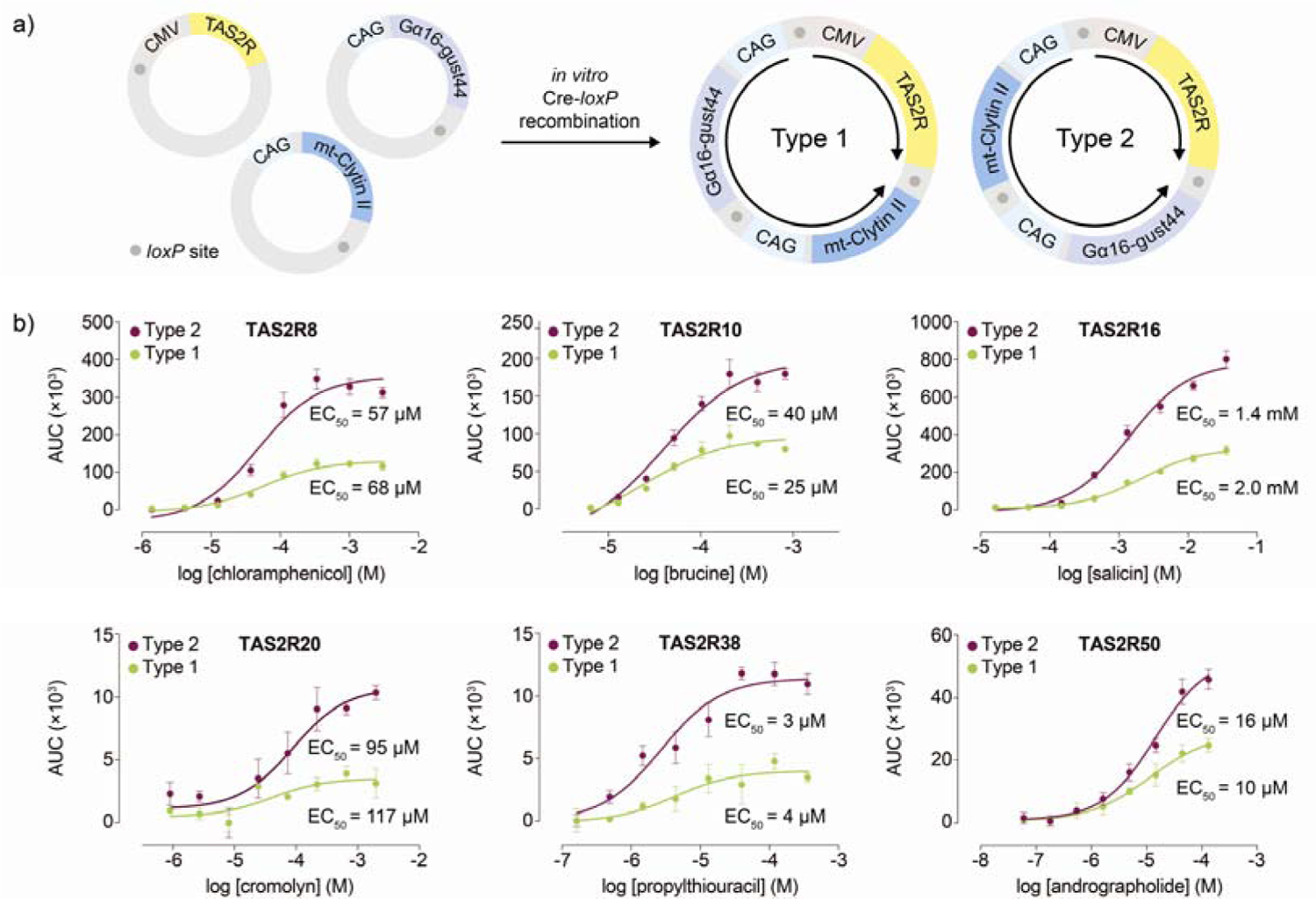
Construction of multigene expression vectors and biochemical analysis. a) Assembly of individual expression vectors encoding TAS2R, mt-clytin II, and Gα gust44 by *in vitro* Cre-*loxP* recombination to generate type 1 and type 2 multigene vectors. With TAS2R as the gene of reference, the expression vectors are arranged in the following sequences in clockwise fashion: TAS2R–mt-clytin II–Gα16-gust44\ (type 1) and TAS2R–G 16-gust44–mt-clytin II (type 2). In both type 1 and type 2 plasmids, TAS2R is divergently orientated from mt-clytin II and Gα16-gust44 chimera. Black arrows indicate the direction of expression from the CMV or CAG promoter. Grey circles represent *loxP* sites. Only relevant elements of the expression constructs are shown for figure clarity. b) Concentration-response curves of TAS2Rs upon stimulation with their cognate agonists in the bioluminescence-based intracellular calcium release assay. Type 2 vectors consistently produced larger assay spans than that of type 1 vectors. Data points are shown as mean ± s.e.m. from a representative experiment out of three independent biological replicates performed in technical quadruplicates. EC_50_, half-maximal effective concentration.

Studies of gene positional and orientation effects on protein expression levels in systems using multigene vectors have reported conflicting results. Kriz *et al.*, for instance, observed similar expression levels of three proteins encoded in multigene vectors with different sequence assembly patterns.^15^ In contrast, Underhill *et al.*, reported imbalanced protein expression ratios in a dual-gene, single-plasmid system, with higher expression biased towards the gene that is placed downstream of the first.^16^ These opposing observations have potentially significant implications for the performance of the bioluminescence-based assay. Therefore, the optimal type of multigene construct to utilize for the assay needs to be empirically determined. Type 1 and type 2 multigene vectors of six TAS2Rs (TAS2R8/10/16/20/38/50) were tested against their cognate agonists. Similar EC_50_ values were generated from both type 1 and type 2 plasmids. However, the observation that type 2 plasmids consistently produced overall larger magnitude of responses compared to type 1 plasmids suggests that the former induced an overall higher gene expression (Figure 1b). Henceforth, type 2 multigene constructs were generated for other TAS2Rs and used throughout the study. A variety of potential mechanisms for transcriptional interference arising from the positional effects of genes in the type 1 plasmid could account for its lower gene expression levels, including promoter occlusion, competition for transcription machinery, or creation of antisense RNA that resulted in translational downregulation.^16^

### Bioluminescence-based intracellular calcium release assay generated larger assay spans than that of a fluorescence-based assay

The performance difference between the bioluminescence-based and the fluorescence-based assay was evaluated. Using TAS2R50, agonist-induced calcium responses were examined in both assay modalities using the multigene constructs. The kinetics of both bioluminescence and fluorescence intensities showed ligand concentration dependence, and were characterized by an initial rise in light intensity after agonist addition, followed by an eventual decrease (Figure 2). The kinetics of signal decline in the fluorescence-based assay was prolonged because calcium ions dissociate slowly from the fluorescent dye (Figure 2a). However, bioluminescence signals decayed more rapidly than fluorescence signals due to rapid consumption of the coelenterazine substrate upon calcium binding in mt-clytin II (Figure 2b). Despite this difference, similar EC_50_ values were obtained using both assays by adopting the most appropriate data processing paradigm for each assay (see Methods). A significant advantage of the bioluminescence-based assay is its capability to generate larger assay spans, and therefore deliver higher assay sensitivity and robustness, than fluorescence-based assays (Figure 2). This enhanced sensitivity and large assay window could potentially facilitate the detection and characterization of low potency agonists and partial agonists in screening campaigns with greater confidence. These results indicate that the bioluminescence-based assay is suitable for the detection of TAS2R agonists, and has a markedly higher signal-to-noise ratio (SNR) (defined as mean signal - mean background)/standard deviation of background) (SNR_bioluminescence(AUC)_ = 90 or SNR_bioluminescence((ΔF/F)_ = 30) compared to the fluorescence-based assay (SNR_fluorescence((ΔF/F)_ = 15).

**Figure 2.**
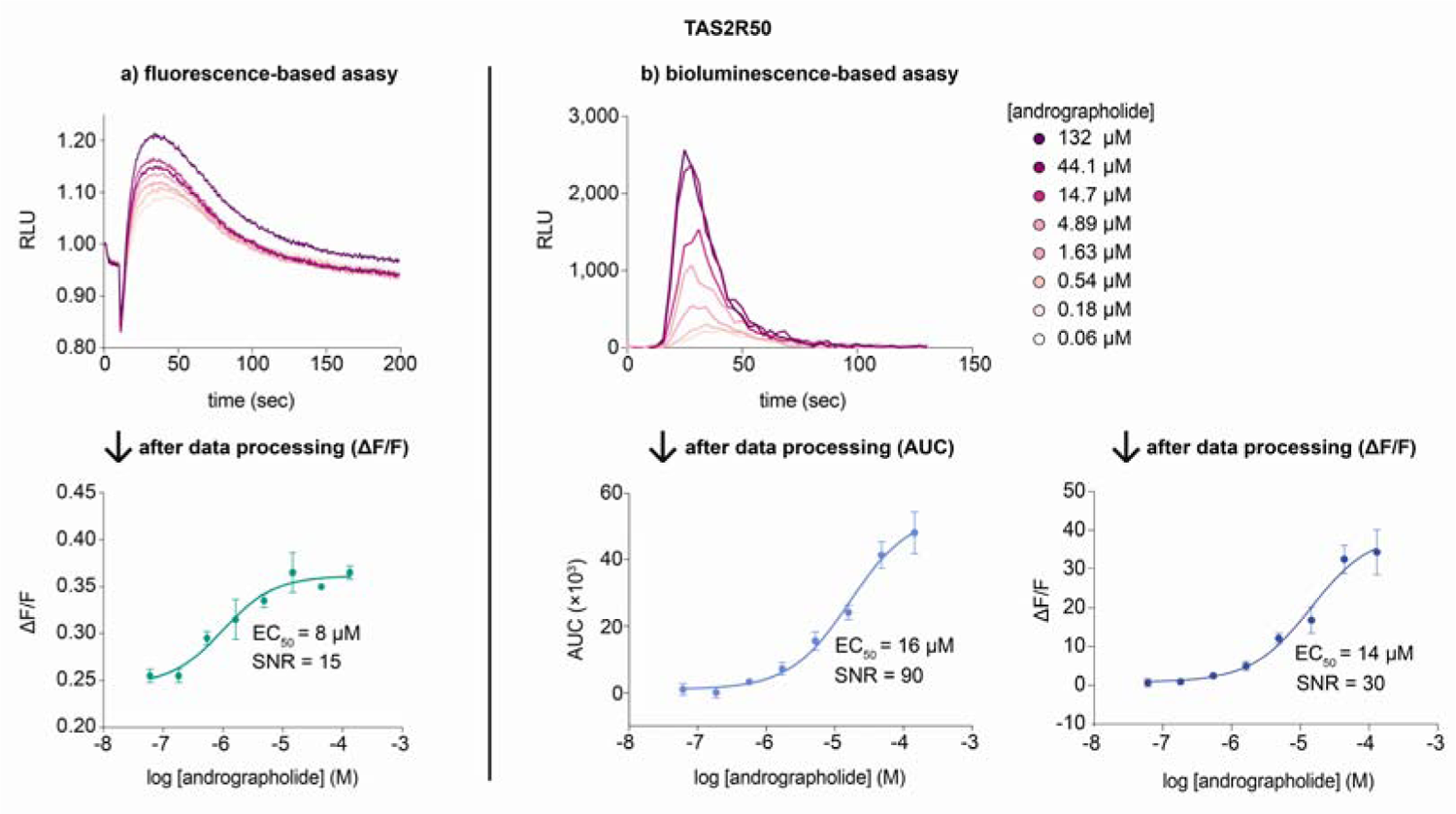
Cells expressing TAS2R50 were loaded with Fluo4 and coelenterazine for the a) fluorescence-based and b) bioluminescence-based assay, respectively, and the emission signals were recorded as a function of time after agonist introduction. Both assay modalities displayed concentration-dependent responses, but the kinetics of signal decay was more rapid for the bioluminescence-based assay compared to the fluorescence-based assay. After data processing of the raw data, the bioluminescence-based assay produced a larger signal-to-noise ratio than that of the fluorescence-based assay. Data points are shown as mean ± s.e.m. from a representative experiment out of at least two independent biological replicates performed in technical quadruplicates. EC_50_, half-maximal effective concentration. SNR, signal-to-noise ratio.

### Validation of the bioluminescence-based TAS2R assay

The bioluminescence-based assay was used to characterize the functional properties of various human TAS2Rs. For each TAS2R, one or more of its cognate agonists were tested against either the AD-293 or 293AD cell line, or both. Both cell lines were used because of their stronger adherent properties, which facilitate media changes on the day of the FLIPR assay. However, because both cell lines are generated differently, some disparities in gene expression capacities due to a myriad of factors could be reasonably expected. Consequently, some TAS2Rs may be well expressed in one cell line but not the other, and this needs to be empirically evaluated. In general, the EC_50_ values obtained from the bioluminescence-based assay were comparable to those reported in the literature (Figure 3, Supplementary Figure 2, and Table 1). However, deviations between our experimental and published potency values for some ligands (i.e. TAS2R4: stevioside; TAS2R7: Mg^2+^, Mn^2+^, Zn^2+^) were observed, which could be attributed to the use of a different assay format, readout and processing parameters, use of different cell lines with resultant potential differences in TAS2R expression levels, and quality and methodological differences in ligand preparations. Because potency values could be obtained that were consistent with the literature, or close to the reported minimal effective stimulatory concentration upon further testing of other TAS2R4/7 agonists, the deviation in EC_50_ values could be largely attributed to qualitative differences in the ligands (Supplementary Figure 3). Calcium responses induced by the tested cognate agonists reflected specific TAS2R stimulation. Non-specific activation of endogenous GPCRs, calcium channels, or cellular effectors was discounted because cells in our control conditions (i.e. transfected to express both Gα16-gust44 and mt-clytin II, or solely mt-clytin II) produced flat, non-dose dependent calcium responses, or responses that resulted in partial dose-response curves, indicating extremely weak potency of the ligands when challenged with TAS2R agonists (Supplementary Table 4 and Supplementary Figure 4).

**Figure 3.**
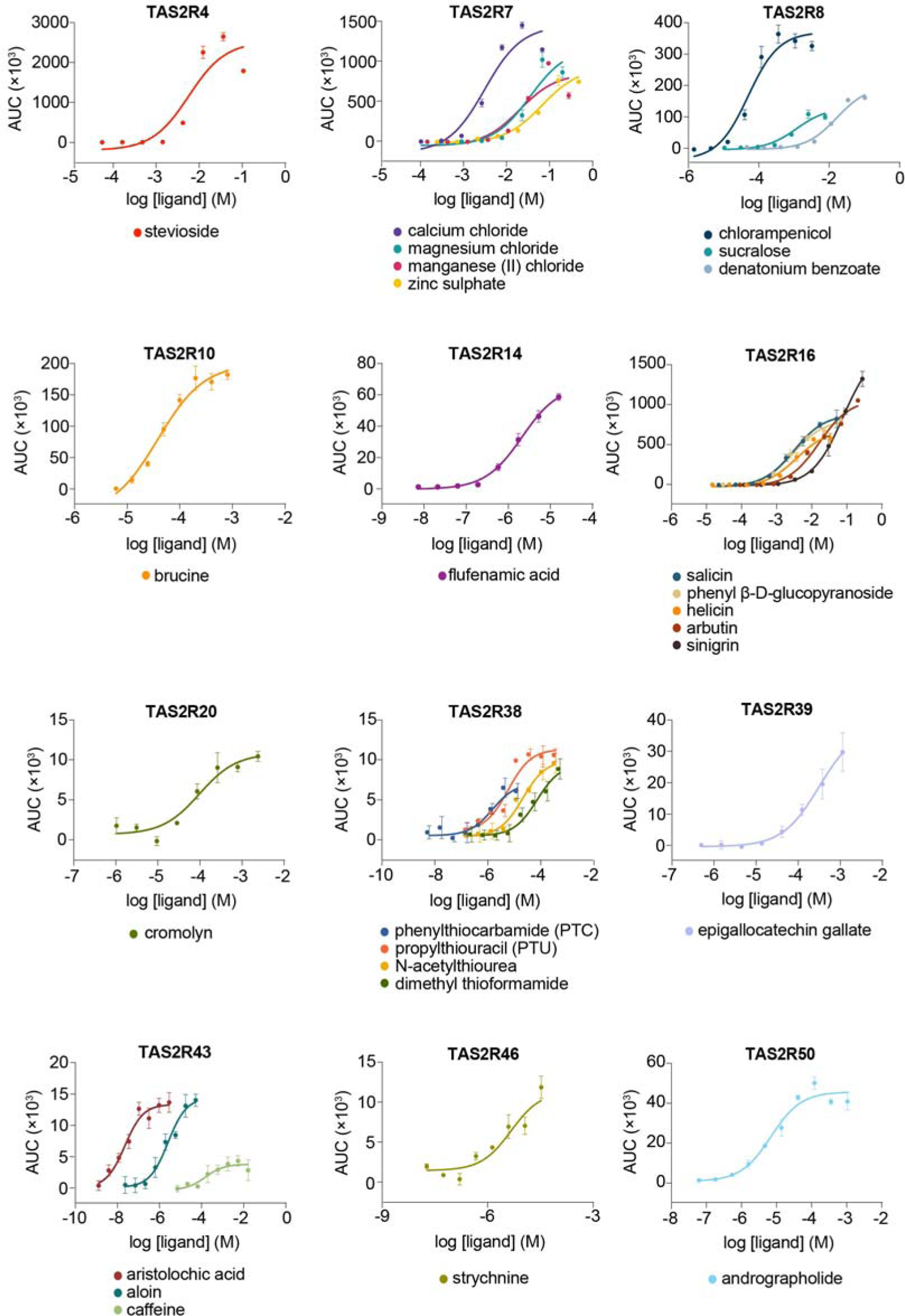
Concentration-response curves of TAS2Rs upon stimulation with their cognate agonists in the bioluminescence-based intracellular calcium release assay in AD-293 cells. Data points are shown as mean ± s.e.m. from a representative experiment out of at least two independent biological replicates performed in at least technical duplicates.

**Table 1.** EC_50_ values of TAS2R agonists. Values indicate the mean ± s.e.m. (n = 2-4).

| Bitter Receptor | Compound | EC<sub>50</sub> values |  | Published EC<sub>50</sub> values |
| --- | --- | --- | --- | --- |
|  |  | 293AD | AD-293 |  |
| TAS2R3 | Chloroquine diphosphate | 45 ± 14 μM | - | 172 ± 29 μM<sup>5</sup> |
| TAS2R4 | Stevioside | 7 ± 2 mM | 7 ± 3 mM | 341 ± 34 μM<sup>17</sup> |
| TAS2R5 | Epigallocatechin gallate | 47 ± 11 μM | - | 12.3 ± 3.63 μM<sup>18</sup> |
| TAS2R7 | Calcium chloride | 5 ± 0.5 mM | 4 ± 1 mM | 5.27 ± 0.5 mM<sup>19</sup> |
| | Magnesium chloride | - | 75 ± 9 mM | 6.07 ± 1.07 mM<sup>19</sup><br>10 ± 19.6 mM<sup>20</sup> |
| | Zinc sulphate | - | 83 ± 11 mM | 33.36 ± 0.14 mM<sup>19</sup> |
| | Manganese (II) chloride | - | 28 ± 5 mM | 6.59 ± 1.73 mM<sup>19</sup><br>10 ± 1.7 mM<sup>20</sup> |
| TAS2R8 | Chloramphenicol | 18 ± 2.7 μM | 70 ± 12 μM | 41 μM<sup>21</sup> |
| | Denatonium benzoate | - | 1 ± 0.1 mM | - |
| | Sucralose | - | 11 ± 4 mM | - |
| TAS2R9 | Pirenzepine | 4 ± 0.4 mM | - | 1.8 mM<sup>22</sup> |
| TAS2R10 | Brucine | 21 ± 6.5 μM | 42 ± 3 μM | - |
| TAS2R13 | Oxyphenonium | 161 ± 16 μM | - | - |
| TAS2R14 | Flufenamic acid | 422 ± 96 nM | 1,490 ± 481 nM | 137 ± 17 nM<sup>5</sup><br>238 ± 12.9 nM<sup>23</sup> |
| | Aristolochic acid | 2.2 ± 1 μM | - | - |
| | Picrotoxinin | 54 ± 8.4 μM | - | 2.6 μM<sup>24</sup><br>13.16 ± 0.93 μM<sup>25</sup><br>18 μM<sup>26</sup> |
| TAS2R16 | Salicin | $0.4 \pm 0.2$ mM | $2 \pm 0.6$ mM | $0.8 \pm 0.2$ mM<sup>27</sup><br>$1.4 \pm 0.2$ mM<sup>5</sup><br>$0.417$ mM<sup>28</sup><br>$0.22$ mM<sup>29</sup> |
| | Helicin | - | $3 \pm 0.2$ mM | $2.3 \pm 0.4$ mM<sup>5,30</sup> |
| | Arbutin | - | $5 \pm 1.2$ mM | $5.5 \pm 1.9$ mM<sup>27</sup><br>$5.8 \pm 0.9$ mM<sup>5</sup><br>$1.34$ mM<sup>29</sup> |
| | Sinigrin | - | $70 \pm 18$ mM | $0.23$ mM<sup>31</sup> |
| | Phenyl β-D-glucopyranoside | - | $2 \pm 0.1$ mM | $1.1 \pm 0.1$ mM<sup>30</sup><br>$0.38$ mM<sup>29</sup> |
| TAS2R20 | Cromolyn | $35 \pm 7$ μM | $73 \pm 26$ μM | $45 \pm 25$ μM<sup>5</sup><br>$64.37 \pm 13$ μM<sup>32</sup> |
| TAS2R30 | Amarogentin | $3 \pm 1$ μM | - | - |
| TAS2R31 | Aristolochic acid | $186 \pm 70$ nM | - | $455 \pm 5.3$ nM<sup>5</sup><br>$130 \pm 10$ nM<sup>33</sup><br>$240$ nM (WMVI)<sup>34</sup><br>$810$ nM (RLAV)<sup>34</sup> |
| TAS2R38 | Propylthiouracil | $0.9 \pm 0.3$ μM | $6 \pm 2$ μM | $2.1 \pm 0.9$ μM<sup>5</sup><br>$1.5$ μM<sup>35</sup><br>$2.2$ μM<sup>36</sup> |
| | Phenylthiocarbamide | - | $2 \pm 0.4$ μM | $1.1 \pm 0.5$ μM<sup>5</sup><br>$6$ μM<sup>37</sup><br>$4.5$ μM<sup>35</sup><br>$2.3$ μM<sup>36</sup> |
| | N-acetylthiourea | - | $16 \pm 8$ μM | $25 \pm 16$ μM<sup>5</sup> |
| | Dimethyl thioformamide | - | $79 \pm 12$ μM | $59 \pm 17$ μM<sup>5</sup> |
| TAS2R39 | Epigallocatechin gallate | $141 \pm 44 \mu\text{M}$ | $362 \pm 81 \mu\text{M}$ | $8.50 \pm 2.84 \mu\text{M}^{18}$<br>$161 \mu\text{M}^{38}$<br>$181.6 \mu\text{M}^{39}$ |
| TAS2R43 | Aristolochic acid | $20 \pm 4 \text{ nM}$ | $26 \pm 3 \text{ nM}$ | $8 \text{ nM}^{34}$<br>$81 \pm 0.8 \text{ nM}^5$ |
| | Aloin | - | $5 \pm 3 \mu\text{M}$ | $1.2 \mu\text{M}^{34}$<br>$2.8 \pm 0.4 \mu\text{M}^5$<br>$35 \mu\text{M}^{36}$ |
| | Caffeine | - | $0.39 \pm 0.12 \text{ mM}$ | $0.94 \pm 0.14 \text{ mM}^{40}$ |
| TAS2R46 | Strychnine | $309 \pm 73 \text{ nM}$ | $1,201 \pm 433 \text{ nM}$ | $0.39 \pm 0.08 \mu\text{M}^{41}$<br>$0.43 \pm 0.02 \mu\text{M}^{42}$<br>$3.47 \mu\text{M}^{43}$ |
| TAS2R50 | Andrographolide | $2 \pm 0.3 \mu\text{M}$ | $16 \pm 3 \mu\text{M}$ | $22.9 \pm 4.9 \mu\text{M}^5$ |

The data obtained from the bioluminescence-based assay appeared to have resulted in elevated apparent potencies, and in many cases the measurements from published fluorescence-based assays seemed to be more potent (Table 1). However, careful consideration is warranted before making direct comparisons of EC_50_ values between different assays. The kinetics of response to calcium binding and subsequent calcium dissociation from a fluorophore is significantly different from that of a photoprotein and will result in divergent effects on time to peak and rate of signal decrease. For this reason, different methods were used to transform kinetic bioluminescence and fluorescence data into sigmoidal dose-response curves, resulting in the derivation of different potency values between the two assay formats. Furthermore, experimental settings used in the assay (e.g. dispensing speed and height) can exert profound effects on the response kinetics, and consequently the determination of EC_50_ values. Altogether, the real significance of potency or efficacy values determined from a particular experimental format must be acknowledged before comparing them with values from different assays.

Furthermore, the assay can be used to evaluate the E_max_ of various ligands and classify ligands as full agonists or partial agonists. For instance, in the concentration response curve for TAS2R43, both aristolochic acid and aloin could be considered full agonists, with the former being more potent than the latter. However, caffeine did not engender a 100% response level as that of aristolochic acid and aloin even at the highest concentration tested, thereby indicating its partial agonist activity towards TAS2R43 (Figure 3).

### The bioluminescence-based assay enables evaluation of TAS2R agonists within an autofluorescent matrix

The generation of bioluminescent signals does not require an excitation light source and is free from interference by autofluorescence, potentially allowing complex biological samples that contain autofluorescent compounds to be screened. To evaluate the possibility of screening autofluorescent samples in the bioluminescence-based assay, agonist solutions were prepared by dilution in Luria-Bertani (LB) broth, a complex medium that displays autofluorescence when excited with blue light due to the presence of compounds such as flavin, and tested against TASR50 in both the fluorescence- and bioluminescence-based assays. In the fluorescence-based assay, concentration-dependent cellular responses could only be observed when andrographolide was dissolved in a non-autofluorescent medium like Hanks’ Balanced Salt Solution (HBSS) (Figure 4a). The use of andrographolide dissolved in LB medium generated uniformly high fluorescent signals, effectively masked the cellular responses to the cognate agonists, and prevented the generation of dose-response curves (Figure 4a). In contrast, dose-dependent cellular responses of andrographolide prepared in LB medium could be detected in the bioluminescence-based assay, and the EC_50_ value obtained was in good agreement with that obtained from testing andrographolide in dissolved in HBSS (Figure 4b and Table 1). Interestingly, the signal-to-noise ratio of the assay is higher in the complex LB medium compared to HBSS buffer. We compared LB medium and HBSS in terms of their activation of calcium responses in two control settings (i.e. cells transfected to express both Gα16-gust44 and mt-clytin II, or solely mt-clytin II). In both settings, LB medium alone clearly activated higher calcium responses than HBSS (Supplementary Figure 5). Therefore, the application of andrographolide dissolved in LB medium added a signal from TAS2R50 activation on top of LB-broth induced signals to result in a higher signal-to-noise ratio.

**Figure 4.**
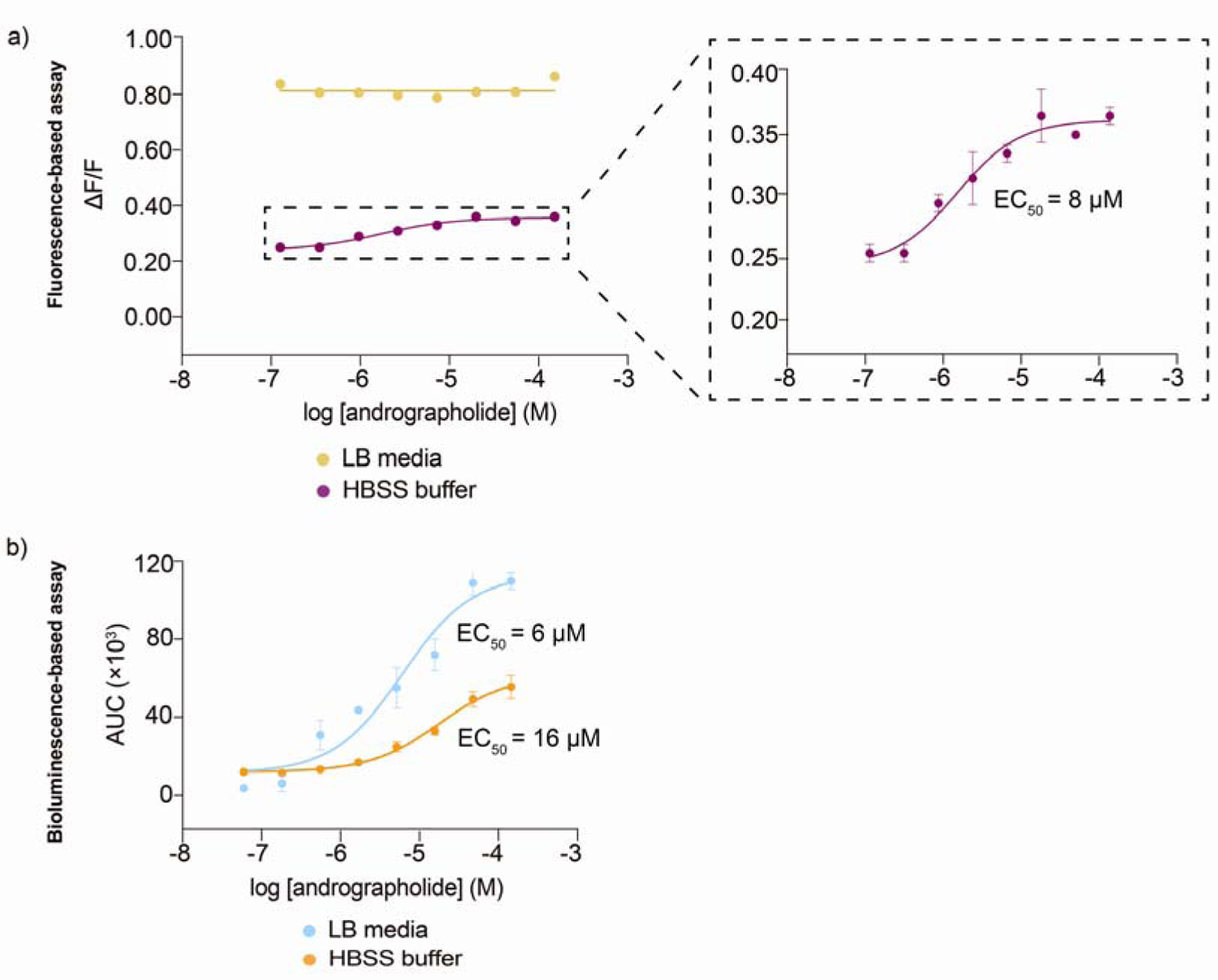
Dose-response curve of TAS2R50 in response to andrographolide stimulation when dissolved in different media. a) Andrographolide dissolved in LB medium produced uniformly high fluorescence signals. When andrographolide was dissolved in HBSS buffer, a clear dose-response curve could be discerned, with an inset showing an enlarged view of the functional response curve. b) Concentration-response curve of TAS2R50 upon stimulation by andrographolide dissolved in LB and HBSS buffer obtained from the bioluminescence-based assay. Data points are shown as mean ± s.e.m. from a representative experiment out of at least two independent biological replicates performed in technical quadruplicates. EC_50_, half-maximal effective concentration.

### Screening of signal sequences to enhance functional bioluminescent readout of TAS2Rs

The observation that the SST_3_ signal sequence produced different assay spans for different TAS2Rs in the bioluminescence-based assay suggested that its ability to mediate receptor surface expression varied with the identity of the TAS2R. Given that GPCRs constitute a vast protein family, we speculate that some TAS2Rs could exhibit higher cell surface expression when grafted with putative signal sequences of other GPCRs to produce larger assay spans in the bioluminescence-based calcium assay. Hence, a cell surface expression assay was developed based on the NanoBiT technology. TAS2R constructs incorporating an N-terminal signal sequence followed by a HiBiT tag, an 11-amino acid tag (VSGWRLFKKIS), were generated and transfected into AD-293. Plasma-membrane localized TAS2Rs will display the HiBiT tag extracellularly, enabling it to form a highly luminescent enzyme when incubated with a non-lytic reagent containing LgBiT protein, which binds with high affinity (K_D_ = 0.7 nM) to the HiBiT tag, and furimazine substrate.^44^ The luminescence level generated after addition of the reagent is directly proportional to the cell surface expression level of the HiBiT-tagged TAS2R. Accordingly, signal sequences that are more effective at promoting TAS2R translocation to the plasma membrane will generate higher luminescence signals (Figure 5a).

**Figure 5.**
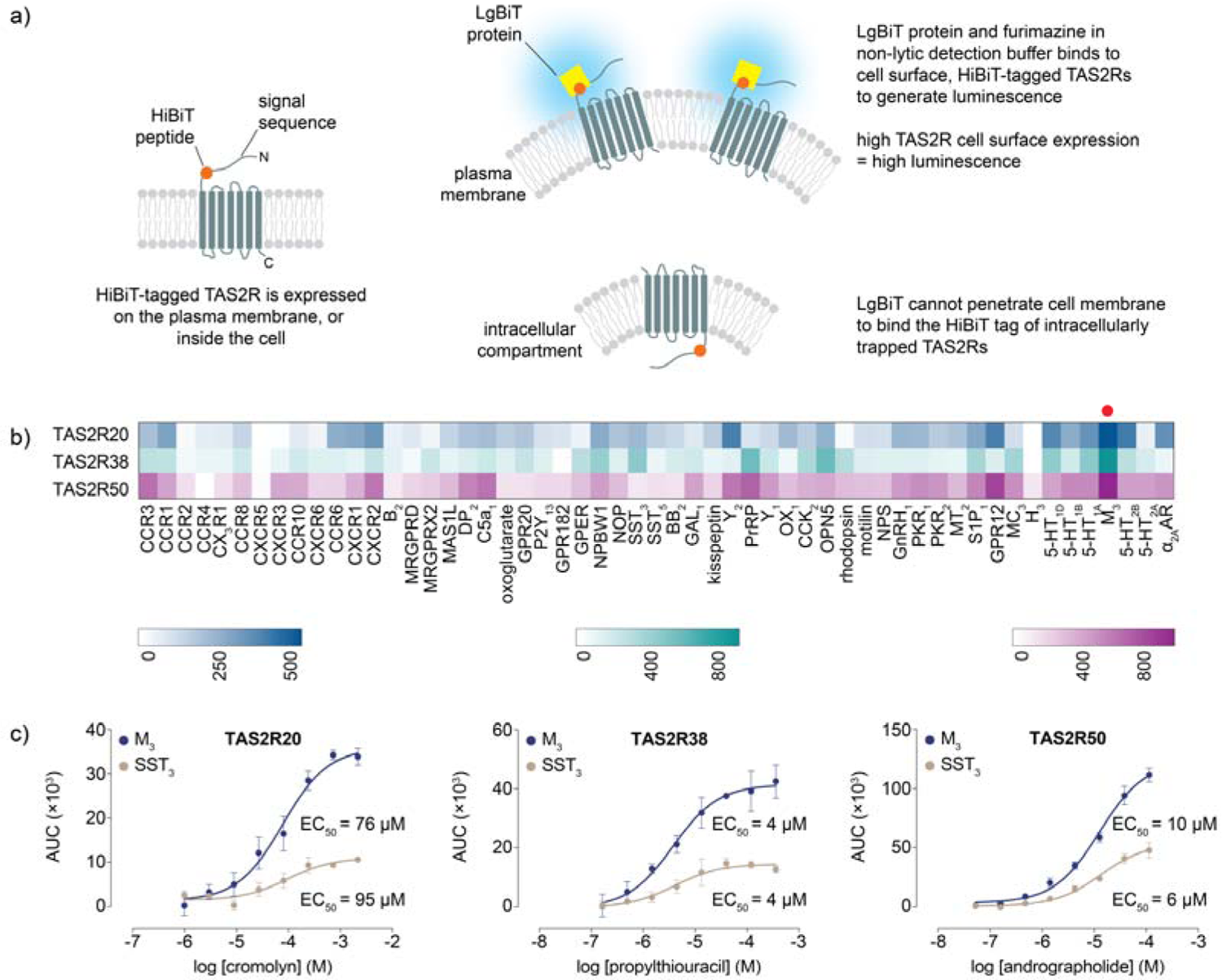
Screening of signal sequences to increase functional expression of TAS2R20/38/50 at the plasma membrane. a) Schematic of the design and principles of the cell surface expression HiBiT assay. b) Heat map of the fold increase in luminescence signals of TAS2R20/38/50 tagged with various N-terminal signal sequences compared with untransfected cells. The signal sequence derived from the M_3_ receptor, indicated by a red circle, generated the highest surface TAS2R expression for TAS2R20/38/50. c) Concentration-response curves of TAS2R20/38/50 upon stimulation with their cognate agonists in the bioluminescence-based intracellular calcium release assay. M_3_-TAS2Rs had higher overall magnitude of responses compared to SST_3_-TAS2Rs. The identities of GPCRs used for screening are abbreviated according to the nomenclature employed by the IUPHAR/BPS Concise Guide to Pharmacology (http://www.guidetopharmacology.org). Adrenergic receptor is abbreviated to AR for simplicity. Data points are shown as mean ± s.e.m. from a representative experiment out of three independent biological replicates performed in technical quadruplicates. EC_50_, half-maximal effective concentration.

Putative signal sequences from 55 randomly selected, non-olfactory Class A GPCRs, which share up to 29% sequence homology with TAS2Rs in the transmembrane domains, were tested for their effect on cell surface expression of HiBiT-tagged TAS2Rs.^3^ This approach was applied to three TAS2Rs (TAS2R20/38/50) that showed relatively small signal spans in the bioluminescence-based intracellular calcium release assay. Results from the cell surface expression assay showed that the signal sequence of muscarinic acetylcholine M3 receptor (M_3_) caused the highest degree of plasma membrane translocation of TAS2R20/38/50 (Figure 5b). Consequently, type 2 multigene vectors containing the respective M_3_-TAS2Rs (without HiBiT tag) were generated and profiled using their cognate agonists to determine whether increased receptor trafficking to the plasma membrane translated to augmented signaling responses in the bioluminescence-based calcium assay. Concentration-response relationships revealed similar EC_50_ values were obtained using both M_3_-TAS2Rs and SST_3_-TAS2Rs, indicating that N-terminal alteration of TAS2Rs did not affect receptor-ligand interactions and functional responses. More importantly, the dose-response curves demonstrated overall more pronounced response magnitudes, with a 2- to 3-fold increase in signal window being observed, from using M_3_-TAS2Rs than SST_3_-TAS2Rs, suggesting that the M_3_ sequence facilitated greater export of functional TAS2Rs to the cell surface than the SST_3_ tag (Figure 5c).

Various studies have highlighted the role of signal sequence *N*-glycosylation in the trafficking of GPCRs to the cell surface. To probe the importance of glycosylation in mediating the transport of TAS2Rs to the cell surface, asparagine residues within the M_3_ and SST_3_ signal sequences (without HiBiT tag) were mutated to alanine in order to probe the effect of signal sequence *N*-glycosylation on functional expression of TAS2R20/38/50 in the bioluminescence-based assay. Receptors TAS2R20/38/50 which were grafted with *N*-glycosylation-deficient signal sequences [M_3_(Asn Ala)-TAS2R and SST_3_(Asn Ala)-TAS2R] did not evoke cellular responses upon application of their cognate agonists, suggesting a failure of receptor transport to the plasma membrane and intracellular trapping of these TAS2Rs (Figure 6).

**Figure 6.**
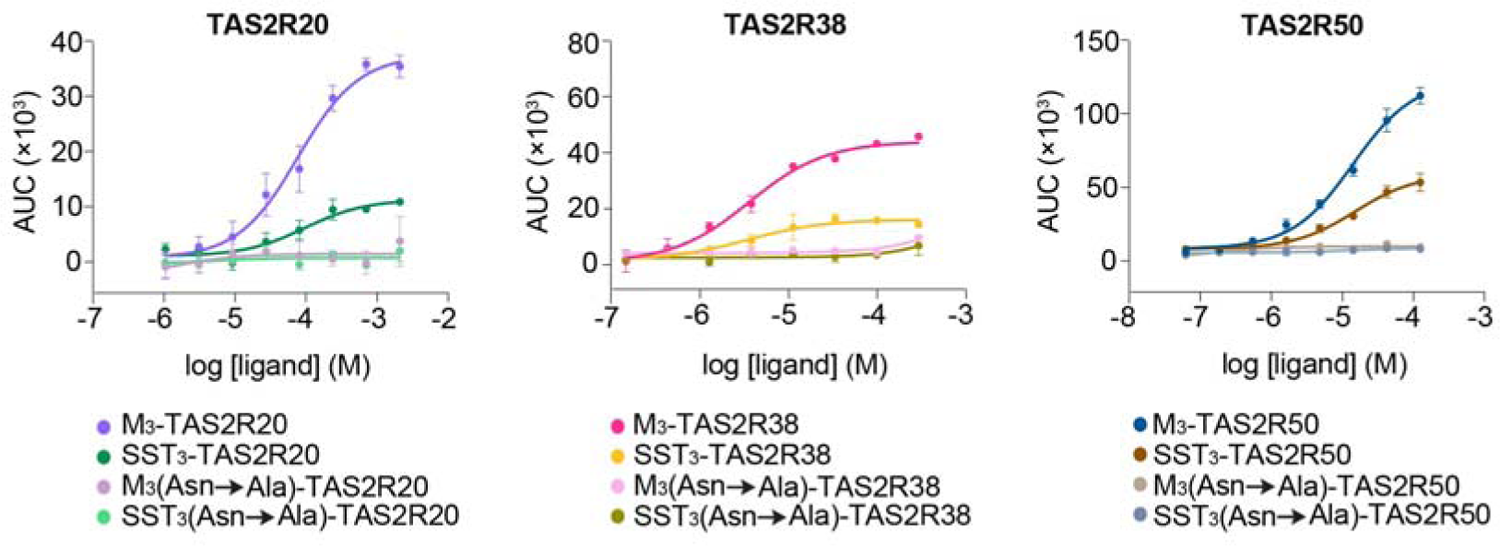
Effect of *N*-glycosylation competent or deficient M_3_ and SST_3_ signal sequences on the functional responses of correspondingly tagged TAS2R20/38/50 upon stimulation by their cognate agonists. TAS2R20/38/50 grafted with *N*-glycosylation-deficient M_3_ and SST_3_ signal sequences were resistant to activation by their cognate agonists. Data points are shown as mean ± s.e.m. from a representative experiment out of three independent biological replicates performed in technical quadruplicates.

## Discussion

Heterologously expressed TAS2Rs suffer from poor cell surface expression as non-native cells may be deficient in the appropriate cofactors or trafficking apparatus for folding and exporting TAS2Rs to the plasma membrane. Poor localization of TAS2Rs to the cell surface in heterologous systems is associated with diminished receptor signaling output and functionality. One of the more commonly used strategies to enhance the density of TAS2Rs at the cell surface involves fusing an N-terminal signal sequence of either the SST_3_, or Rho receptor to the TAS2R. We adopted this approach, specifically co-expressing SST_3_-TAS2R with Gαclytin II, to develop and validate a bioluminescence-based TAS2R intracellular calcium release assay. The bioluminescence-based TAS2R assay produced greater assay spans than that of fluorescence-based assays. This is largely attributed to the utilization of mt-clytin II as the calcium reporter. Calcium-binding photoproteins in general are useful reporters for studying calcium-mediated signaling processes. The mt-clytin II-coelenterazine complex displays negligible basal luminescence, and responds almost immediately to calcium binding by emitting a flash luminescence, followed by an exponential decay due to consumption of the coelenterazine substrate.^10^ Its fast calcium binding and dissociation kinetics render it a suitable indicator to report on swift, transient changes in calcium ion concentrations and study rapid receptor-ligand interactions. Mitochondrial targeting of clytin II is crucial to generating higher luminescence intensity than that of cytosol-localized clytin II, as inositol 1,4,5-triphosphate (IP_3_)-induced calcium release from the ER subsequent to ligand activation is thought to result in the formation of ER-proximal, calcium-rich microdomains that enables the closely apposed mitochondria to sense a higher initial rise in calcium concentration than would be experienced by the bulk cytosol.^14^ Furthermore, while other calcium-sensitive photoproteins could be employed, mt-clytin II was chosen as it has been shown to generate higher luminescence intensity and a more favorable signal-to-noise ratio in the assay than that of mt-aequorin, mt-clytin, and mt-obelin.^12^

An additional advantage that stems from using a photoprotein reporter is the ability to screen samples that contain autofluorescent substances, which is a common characteristic of plant and food extracts or fractionates. We have demonstrated that the bioluminescence-based assay was able to detect agonist-induced increase in calcium mobilization in a concentration-dependent fashion even when the ligands were dissolved in LB broth, an autofluorescent media. This potentially enables the bioluminescence-based assay to be applied to the screening of plant or food extracts, and could find potential application in the discovery of bitter taste modulators for the food industry.

Another strategy that has been reported to increase the density of TAS2Rs at the plasma membrane involves co-expressing TAS2Rs with accessory factors such as receptor transporting proteins (RTPs), and receptor expression enhancing proteins (REEPs), which have been reported to increase the functional expression of heterologously expressed odorant receptors (ORs) by several probable mechanisms that include assisting in proper folding of ORs, promoting cell surface transport of OR-containing compartments, and serving as coreceptors of ORs by providing appropriate trafficking signals or masking potential OR retention motifs.^45^ However, this method is not widely employed in TAS2R studies probably because the effects of RTPs and REEPs on TAS2R function appeared to be receptor-specific.^46^

As mentioned previously, the more frequently utilized strategy involves tagging TAS2Rs with either the SST_3_ or Rho signal sequence (referred to as SST_3_-TAS2R or Rho-TAS2R). The amplitude of signaling responses in calcium assays was greater by using signal sequence-tagged TAS2Rs than by co-expression of accessory proteins, suggesting that the former elicited higher functional receptor expression at the plasma membrane, and provided a motivation to screen other putative signal sequences from various non-olfactory, class A GPCRs to explore whether this approach can improve assay functional readout in the bioluminescence-based assay by improving TAS2R cell surface expression.^46^ The extracellular N-termini of TAS2Rs are predicted to be generally short, span 1-30 amino acid residues, and are devoid of signal peptide sequences and *N*-glycosylation sites (Supplementary Table 5). However, TAS2Rs are predicted, and some have even been shown, to be predominantly *N*-glycosylated at a conserved and consensus site in the second extracellular loop of all 25 human TAS2Rs (Supplementary Table 5), which has been demonstrated to be important for receptor trafficking to the cell surface.^47^ While the approach of incorporating N-terminal sequences to improve receptor cell surface trafficking has been applied to various GPCRs, we note that hitherto only signal sequences from the SST_3_ and Rho receptors have been employed for TAS2R studies.^48^ We were further interested in exploring the effect of changing the N-terminal signal sequences of TAS2Rs to that of other GPCRs on the functional readout of the bioluminescence-based TAS2R assay. Therefore, we selected 3 TAS2Rs (TAS2R20/38/50), which have relatively low signal windows using SST_3_ tag, as test cases by investigating the effect of attaching various putative signal sequences from 55 Class A, non-olfactory GPCRs on the functional readout. The screen revealed that these 3 receptors coincidentally had higher overall magnitude of responses when tagged with the signal sequence from the M_3_ receptor.

*N*-glycosylation of GPCRs is a ubiquitous type of post-translational modification that performs diverse receptor-specific roles, including functions in cell surface expression, protein folding, structural maturation, oligomerization, ligand recognition, and coupling to signal transduction systems.^49, 50^ The SST_3_ and M_3_ export tags contain 2 (N18 and N31) and 5 (N6, N15, N41, N48, and N52) potential *N*-glycosylation sites in their protein sequences, respectively. The ablation of one *N*-glycosylation site on the N-terminus of an OR impeded its translocation to the plasma membrane, suggesting the importance of N-terminal *N*-glycans for proper functioning and localization of chemosensory receptors.^51^ Similar positional effects of *N*-glycans on cell surface expression have also been observed for some Class A GPCRs, such as the bradykinin receptor, and dopamine receptor, whereby disruption of receptor *N*-glycosylation inhibited their translocation to the plasma membrane.^49, 52^ In a heterologous system that lacks potentially specialized folding or trafficking machinery in taste receptor cells, the presence of artificially introduced *N*-glycans may assist in proper TAS2R folding by engaging glycoprotein-dedicated chaperone systems that involve calnexin, calreticulin, and binding immunoglobulin protein (BiP).^47, 53^ In addition, *N*-glycosylation may enhance TAS2R export to the cell surface by favoring receptor accumulation in lipid rafts, or promoting interactions with signaling compartments involved in receptor trafficking.^54^

Compared with the SST_3_ export tag, the introduction of more *N*-glycosylation sites to the protein sequences of TAS2R20/38/50 by using the M_3_ signal sequence may further enhance TAS2R cell surface expression by improving receptor folding, or by better overcoming potential inhibitions on receptor transportation to the plasma membrane imposed by protein motifs, or domains specific to these three TAS2Rs, which we were unable to identify. This implies that the cell surface trafficking of some TAS2Rs in a heterologous cell system may depend on the extent of *N*-glycosylation. In fact, individual TAS2Rs, and even other chemoreceptors, have different requirements for auxiliary factors in relation to cell surface trafficking, suggesting that the ease of functional expression of each TAS2R is linked to its identity or protein sequence, and that some TAS2Rs may face an intrinsically greater obstacle to be expressed and exported. This was exemplified by the recalcitrance of native TAS2R10/38 to ligand activation despite RTP and REEP coexpression, reflecting a failure in receptor surface expression, in contrast to the independence of wild type TAS2R14/43, which exhibited similar agonist-mediated signaling responses regardless of the presence of accessory proteins.^46^ In another study, some TAS2Rs (e.g. TAS2R14/46) have been shown to be functional even when expressed without a signal sequence tag, suggesting that the extent of *N*-glycosylation in the receptor coding sequences influences the level of receptor localization to the cell surface.^47^ TAS2R20/38 and TAS2R50 are predicted to contain 2 and 1 *N*-glycosylation sites respectively in their protein sequences (Supplementary Table 5), suggesting that TAS2R20/38 would be inherently better expressed than TAS2R50 in the absence of an exogenous signal sequence. However, without empirical evidence linking actual number of *N*-glycosylated residues to receptor cell surface expression levels, we could only speculate about whether the approach of introducing more *N*-glycosylation sites into the receptor would indubitably promote cell surface expression for all TAS2Rs. Further studies will be needed to elucidate the mechanisms of the M_3_ signal sequence in inducing greater magnitude of responses, and whether this effect could be extended to other TAS2Rs or GPCRs.

## Acknowledgements

We thank Dr. Ann Koay, Dr. Ng Siew Bee, Dr. Nicholas Lindley, and Dr. Hazel Khoo of the Singapore Institute of Food and Biotechnology Innovation for supporting this project.

