## Supplementary Tables and Figures for "Construction of a bioluminescence-based assay for bitter taste receptors (TAS2Rs)"

**Supplementary Table 1. Amino acid sequences of the SST_3_-TAS2Rs used in this study.**

SST_3_ signal sequence

Linker

Flag tag

| **Bitter taste receptor** | **Amino acid sequence** |
| --- | --- |
| TAS2R3 | MAAVTYPSSVPTTLDPGNASSAWPLDTSLGNASAGTSLAGLAVSGLEMGLTEGVFLILSGTQFTLGILVNCFIELVNGSSWFKTKRMSLSDFIITTLALLRIILLCIILTDSFLIEFSPNTHDSGIIMQIIDVSWTFTNHLSIWLATCLGVLYCLKIASFSHPTFLWLKWRVSRVMVWMLLGALLLSCGSTASLINEFKLYSVFRGIEATRNVTEHFRKKRSEYYLIHVLGTLWYLPPLIVSLASYSLLIFSLGRHTRQMLQNGTSSRDPTTEAHKRAIRIILSFFFLFLLYFLAFLIASFGNFLPKTKMAKMIGEVMTMFYPAGHSFILILGNSKLKQTFVVMLRCESGHLKPGSKGPIFSDYKDDDDK |
| TAS2R4 | MAAVTYPSSVPTTLDPGNASSAWPLDTSLGNASAGTSLAGLAVSGLELRLFYFSAIIASVILNFVGIIMNLFITVVNCKTWVKSHRISSSDRILFSLGITRFLMLGLFLVNTIYFVSSNTERSVYLSAFFVLCFMFLDSSSVWFVTLLNILYCVKITNFQHSVFLLLKRNISPKIPRLLLACVLISAFTTCLYITLSQASPFPELVTTRNNTSFNISEGILSLVVSLVLSSSLQFIINVTSASLLIHSLRRHIQKMQKNATGFWNPQTEAHVGAMKLMVYFLILYIPYSVATLVQYLPFYAGMDMGTKSICLIFATLYSPGHSVLIIITHPKLKTTAKKILCFKKDYKDDDDK |
| TAS2R5 | MAAVTYPSSVPTTLDPGNASSAWPLDTSLGNASAGTSLAGLAVSGLELSAGLGLLMLVAVVEFLIGLIGNGSLVVWSFREWIRKFNWSSYNLIILGLAGCRFLLQWLIILDLSLFPLFQSSRWLRYLSIFWVLVSQASLWFATFLSVFYCKKITTFDRPAYLWLKQRAYNLSLWCLLGYFIINLLLTVQIGLTFYHPPQGNSSIRYPFESWQYLYAFQLNSGSYLPLVVFLVSSGMLIVSLYTHHKKMKVHSAGRRDVRAKAHITALKSLGCFLLLHLVYIMASPFSITSKTYPPDLTSVFIWETLMAAYPSLHSLILIMGIPRVKQTCQKILWKTVCARRCWGPDYKDDDDK |
| TAS2R7 | MAAVTYPSSVPTTLDPGNASSAWPLDTSLGNASAGTSLAGLAVSGLEADKVQTTLLFLAVGEFSVGILGNAFIGLVNCMDWVKKRKIASIDLILTSLAISRICLLCVILLDCFILVLYPDVYATGKEMRIIDFFWTLTNHLSIWFATCLSIYYFFKIGNFFHPLFLWMKWRIDRVISWILLGCVVLSVFISLPATENLNADFRFCVKAKRKTNLTWSCRVNKTQHASTKLFLNLATLLPFCVCLMSFFLLILSLRRHIRRMQLSATGCRDPSTEAHVRALKAVISFLLLFIAYYLSFLIATSSYFMPETELAVIFGESIALIYPSSHSFILILGNNKLRHASLKVIWKVMSILKGRKFQQHKQIDYKDDDDK |
| TAS2R8 | MAAVTYPSSVPTTLDPGNASSAWPLDTSLGNASAGTSLAGLAVSGLEFSPADNIFIILITGEFILGILGNGYIALVNWIDWIKKKKISTVDYILTNLVIARICLISVMVVNGIVIVLNPDVYTKNKQQIVIFTFWTFANYLNMWITTCLNVFYFLKIASSSHPLFLWLKWKIDMVVHWILLGCFAISLLVSLIAAIVLSCDYRFHAIAKHKRNITEMFHVSKIPYFEPLTLFNLFAIVPFIVSLISFFLLVRSLWRHTKQIKLYATGSRDPSTEVHVRAIKTMTSFIFFFFLYYISSILMTFSYLMTKYKLAVEFGEIAAILYPLGHSLILIVLNNKLRQTFVRMLTCRKIACMIDYKDDDDK |
| TAS2R9 (A187) | MAAVTYPSSVPTTLDPGNASSAWPLDTSLGNASAGTSLAGLAVSGLEPSAIEAIYIILIAGELTIGIWGNGFIVLVNCIDWLKRRDISLIDIILISLAISRICLLCVISLDGFFMLLFPGTYGNSVLVSIVNVVWTFANNSSLWFTSCLSIFYLLKIANISHPFFFWLKLKINKVMLAILLGSFLISLIISVPKNDDMWYHLFKVSHEENITWKFKVSKIPGTFKQLTLNLGAMVPFILCLISFFLLLFSLVRHTKQIRLHATGFRDPSTEAHMRAIKAVIIFLLLLIVYYPVFLVMTSSALIPQGKLVLMIGDIVTVIFPSSHSFILIMGNSKLREAFLKMLRFVKCFLRRRKPFVPDYKDDDDK |
| TAS2R10 | MAAVTYPSSVPTTLDPGNASSAWPLDTSLGNASAGTSLAGLAVSGLELRVVEGIFIFVVVSESVFGVLGNGFIGLVNCIDCAKNKLSTIGFILTGLAISRIFLIWIIITDGFIQIFSPNIYASGNLIEYISYFWVIGNQSSMWFATSLSIFYFLKIANFSNYIFLWLKSRTNMVLPFMIVFLLISSLLNFAYIAKILNDYKTKNDTVWDLNMYKSEYFIKQILLNLGVIFFFTLSLITCIFLIISLWRHNRQMQSNVTGLRDSNTEAHVKAMKVLISFIILFILYFIGMAIEISCFTVRENKLLLMFGMTTTAIYPWGHSFILILGNSKLKQASLRVLQQLKCCEKRKNLRVTDYKDDDDK |
| TAS2R13 | MAAVTYPSSVPTTLDPGNASSAWPLDTSLGNASAGTSLAGLAVSGLEESALPSIFTLVIIAEFIIGNLSNGFIVLINCIDWVSKRELSSVDKLLIILAISRIGLIWEILVSWFLALHYLAIFVSGTGLRIMIFSWIVSNHFNLWLATIFSIFYLLKIASFSSPAFLYLKWRVNKVILMILLGTLVFLFLNLIQINMHIKDWLDRYERNTTWNFSMSDFETFSVSVKFTMTMFSLTPFTVAFISFLLLIFSLQKHLQKMQLNYKGHRDPRTKVHTNALKIVISFLLFYASFFLCVLISWISELYQNTVIYMLCETIGVFSPSSHSFLLILGNAKLRQAFLLVAAKVWAKRDYKDDDDK |
| TAS2R14 | MAAVTYPSSVPTTLDPGNASSAWPLDTSLGNASAGTSLAGLAVSGLEGGVIKSIFTFVLIVEFIIGNLGNSFIALVNCIDWVKGRKISSVDRILTALAISRISLVWLIFGSWCVSVFFPALFATEKMFRMLTNIWTVINHFSVWLATGLGTFYFLKIANFSNSIFLYLKWRVKKVVLVLLLVTSVFLFLNIALINIHINASINGYRRNKTCSSDSSNFTRFSSLIVLTSTVFIFIPFTLSLAMFLLLIFSMWKHRKKMQHTVKISGDASTKAHRGVKSVITFFLLYAIFSLSFFISVWTSERLEENLIILSQVMGMAYPSCHSCVLILGNKKLRQASLSVLLWLRYMFKDGEPSGHKEFRESSDYKDDDDK |
| TAS2R16 | MAAVTYPSSVPTTLDPGNASSAWPLDTSLGNASAGTSLAGLAVSGLEIPIQLTVFFMIIYVLESLTIIVQSSLIVAVLGREWLQVRRLMPVDMILISLGISRFCLQWASMLNNFCSYFNLNYVLCNLTITWEFFNILTFWLNSLLTVFYCIKVSSFTHHIFLWLRWRILRLFPWILLGSLMITCVTIIPSAIGNYIQIQLLTMEHLPRNSTVTDKLENFHQYQFQAHTVALVIPFILFLASTIFLMASLTKQIQHHSTGHCNPSMKARFTALRSLAVLFIVFTSYFLTILITIIGTLFDKRCWLWVWEAFVYAFILMHSTSLMLSSPTLKRILKGKCDYKDDDDK |
| TAS2R20 | MAAVTYPSSVPTTLDPGNASSAWPLDTSLGNASAGTSLAGLAVSGLEMSFLHIVFSILVVVAFILGNFANGFIALINFIAWVKRQKISSADQIIAALAVSRVGLLWVILLHWYSTVLNPTSSNLKVIIFISNAWAVTNHFSIWLATSLSIFYLLKIVNFSRLIFHHLKRKAKSVVLVIVLGSLFFLVCHLVMKHTYINVWTEECEGNVTWKIKLRNAMHLSNLTVAMLANLIPFTLTLISFLLLIYSLCKHLKKMQLHGKGSQDPSTKIHIKALQTVTSFLILLAIYFLCLIISFWNFKMRPKEIVLMLCQAFGIIYPSFHSFILIWGNKTLKQTFLSVLWQVTCWAKGQNQSTPDYKDDDDK |
| TAS2R30 | MAAVTYPSSVPTTLDPGNASSAWPLDTSLGNASAGTSLAGLAVSGLEITFLPIIFSILIVVIFVIGNFANGFIALVNSIEWVKRQKISFVDQILTALAVSRVGLLWVLLLHWYATQLNPAFYSVEVRITAYNVWAVTNHFSSWLATSLSMFYLLRIANFSNLIFLRIKRRVKSVVLVILLGPLLFLVCHLFVINMDETVWTKEYEGNVTWKIKLRSAMYHSNMTLTMLANFVPLTLTLISFLLLICSLCKHLKKMQLHGKGSQDPSTKVHIKALQTVTSFLLLCAIYFLSMIISVCNFGRLEKQPVFMFCQAIIFSYPSTHPFILILGNKKLKQIFLSVLRHVRYWVKDRSLRLHRFTRGALCVFDYKDDDDK |
| TAS2R31 (WMVI) | MAAVTYPSSVPTTLDPGNASSAWPLDTSLGNASAGTSLAGLAVSGLETTFIPIIFSSVVVVLFVIGNFANGFIALVNSIEWVKRQKISFADQILTALAVSRVGLLWVLLLNWYSTVFNPAFYSVEVRTTAYNVWAVTGHFSNWLATSLSIFYLLKIANFSNLIFLHLKRRVKSVILVMLLGPLLFLACQLFVINMKEIVRTKEYEGNMTWKIKLRSAVYLSDATVTTLGNLVPFTLTLLCFLLLICSLCKHLKKMQLHGKGSQDPSTKVHIKVLQTVIFFLLLCAIYFLSIMISVWSFGSLENKPVFMFCKAIRFSYPSIHPFILIWGNKKLKQTFLSVLRQVRYWVKGEKPSSPDYKDDDDK |
| TAS2R38 (PAV) | MAAVTYPSSVPTTLDPGNASSAWPLDTSLGNASAGTSLAGLAVSGLELTLTRIRTVSYEVRSTFLFISVLEFAVGFLTNAFVFLVNFWDVVKRQPLSNSDCVLLCLSISRLFLHGLLFLSAIQLTHFQKLSEPLNHSYQAIIMLWMIANQANLWLAACLSLLYCSKLIRFSHTFLICLASWVSRKISQMLLGIILCSCICTVLCVWCFFSRPHFTVTTVLFMNNNTRLNWQNKDLNLFYSFLFCYLWSVPPFLLFLVSSGMLTVSLGRHMRTMKVYTRNSRDPSLEAHIKALKSLVSFFCFFVISSCAAFISVPLLILWRDKIGVMVCVGIMAACPSGHAAVLISGNAKLRRAVMTILLWAQSSLKVRADHKADSRTLCDYKDDDDK |
| TAS2R39 | MAAVTYPSSVPTTLDPGNASSAWPLDTSLGNASAGTSLAGLAVSGLELGRCFPPDTKEKQQLRMTKLCDPAESELSPFLITLILAVLLAEYLIGIIANGFIMAIHAAEWVQNKAVSTSGRILVFLSVSRIALQSLMMLEITISSTSLSFYSEDAVYYAFKISFIFLNFCSLWFAAWLSFFYFVKIANFSYPLFLKLRWRITGLIPWLLWLSVFISFSHSMFCINICTVYCNNSFPIHSSNSTKKTYLSEINVVGLAFFFNLGIVTPLIMFILTATLLILSLKRHTLHMGSNATGSNDPSMEAHMGAIKAISYFLILYIFNAVALFIYLSNMFDINSLWNNLCQIIMAAYPASHSILLIQDNPGLRRAWKRLQLRLHLYPKEWTLDYKDDDDK |
| TAS2R43 | MAAVTYPSSVPTTLDPGNASSAWPLDTSLGNASAGTSLAGLAVSGLEITFLPIIFSSLVVVTFVIGNFANGFIALVNSIEWFKRQKISFADQILTALAVSRVGLLWVLLLNWYSTVLNPAFNSVEVRTTAYNIWAVINHFSNWLATTLSIFYLLKIANFSNFIFLHLKRRVKSVILVMLLGPLLFLACHLFVINMNEIVRTKEFEGNMTWKIKLKSAMYFSNMTVTMVANLVPFTLTLLSFMLLICSLCKHLKKMQLHGKGSQDPSTKVHIKALQTVISFLLLCAIYFLSIMISVWSFGSLENKPVFMFCKAIRFSYPSIHPFILIWGNKKLKQTFLSVFWQMRYWVKGEKTSSPDYKDDDDK |
| TAS2R46 | MAAVTYPSSVPTTLDPGNASSAWPLDTSLGNASAGTSLAGLAVSGLEITFLPIIFSILIVVTFVIGNFANGFIALVNSIEWFKRQKISFADQILTALAVSRVGLLWVLVLNWYATELNPAFNSIEVRITAYNVWAVINHFSNWLATSLSIFYLLKIANFSNLIFLHLKRRVKSVVLVILLGPLLFLVCHLFVINMNQIIWTKEYEGNMTWKIKLRSAMYLSNTTVTILANLVPFTLTLISFLLLICSLCKHLKKMQLHGKGSQDPSMKVHIKALQTVTSFLLLCAIYFLSIIMSVWSFESLENKPVFMFCEAIAFSYPSTHPFILIWGNKKLKQTFLSVLWHVRYWVKGEKPSSSDYKDDDDK |
| TAS2R50 | MAAVTYPSSVPTTLDPGNASSAWPLDTSLGNASAGTSLAGLAVSGLEITFLYIFFSILIMVLFVLGNFANGFIALVNFIDWVKRKKISSADQILTALAVSRIGLLWALLLNWYLTVLNPAFYSVELRITSYNAWVVTNHFSMWLAANLSIFYLLKIANFSNLLFLHLKRRVRSVILVILLGTLIFLVCHLLVANMDESMWAEEYEGNMTGKMKLRNTVHLSYLTVTTLWSFIPFTLSLISFLMLICSLCKHLKKMQLHGEGSQDLSTKVHIKALQTLISFLLLCAIFFLFLIVSVWSPRRLRNDPVVMVSKAVGNIYLAFDSFILIWRTKKLKHTFLLILCQIRCDYKDDDDK |

**Supplementary Table 2.** Putative signal sequences of 55 non-olfactory Class A GPCRs used in the study.

| **Gene name** | **Protein name** | **Receptor** | **Putative signal sequence** |
| --- | --- | --- | --- |
| ADRA2A | Alpha-2A adrenergic receptor | α_2A_ | MGSLQPDAGNASWNGTEAPGGGARATPYSLQVT |
| BDKRB2 | B2 bradykinin receptor | B_2_ | MLNITSQVLAPALNGSVSQSSGCPNTEWSGWLNVIQ |
| C5AR1 | C5a anaphylatoxin chemotactic receptor 1 | C5a_1_ | MDSFNYTTPDYGHYDDKDTLDLNTPVDKTSNT |
| CCKBR | Gastrin/cholecystokinin type B receptor | CCK_2_ | MDLLKLNRSLQGPGPGSGSSLCRPGVSLLNSSSAGNLSCETPRIRGTGTRELELTIR |
| CCR1 | C-C chemokine receptor type 1 | CCR1 | METPNTTEDYDTTTEFDYGDATPCQKVNERAFGA |
| CCR10 | C-C chemokine receptor type 10 | CCR10 | MGTEATEQVSWGHYSGDEEDAYSAEPLPELCYKADVQAFSRAFQPSVSLTVA |
| CCR2 | C-C chemokine receptor type 2 | CCR2 | MLSTSRSRFIRNTNESGEEVTTFFDYDYGAPCH |
| CCR3 | C-C chemokine receptor type 3 | CCR3 | MTTSLDTVETFGTTSYYDDVGLLCEKADTRALMA |
| CCR4 | C-C chemokine receptor type 4 | CCR4 | MNPTDIADTTLDESIYSNYYLYESIPKPCTKEGIKAF |
| CCR6 | C-C chemokine receptor type 6 | CCR6 | MSGESMNFSDVFDSSEDYFVSVNTSYYSVDSEMLLCSLQEVRQFSRL |
| CCR8 | C-C chemokine receptor type 8 | CCR8 | MDYTLDLSVTTVTDYYYPDIFSSPCDAELIQTNGK |
| CHRM3 | Muscarinic acetylcholine receptor M3 | M_3_ | MTLHSNSTTSPLFPNISSSWVHSPSEAGLPLGTVTQLGSYNISQETGNFSSNDTSSDPLG |
| CX3CR1 | CX3C chemokine receptor 1 | CX_3_CR1 | MSTSFPELDLENFEYDDSAEACYLGDIVAFGT |
| CXCR1 | C-X-C chemokine receptor type 1 | CXCR1 | MSNITDPQMWDFDDLNFTGMPPADEDYSPCMLETETLNK |
| CXCR2 | C-X-C chemokine receptor type 2 | CXCR2 | MEDFNMESDSFEDFWKGEDLSNYSYSSTLPPFLLDAAPCEPESLEINK |
| CXCR3 | C-X-C chemokine receptor type 3 | CXCR3 | MVLEVSDHQVLNDAEVAALLENFSSSYDYGENESDSCCTSPPCPQDFSLNFDR |
| CXCR5 | C-X-C chemokine receptor type 5 | CXCR5 | MNYPLTLEMDLENLEDLFWELDRLDNYNDTSLVENHLCPATEGPLMASFKAVFV |
| CXCR6 | C-X-C chemokine receptor type 6 | CXCR6 | MAEHDYHEDYGFSSFNDSSQEEHQDFLQFSKV |
| GALR1 | Galanin receptor type 1 | GAL_1_ | MELAVGNLSEGNASWPEPPAPEPGPLFGIGVENFVT |
| GNRHR | Gonadotropin-releasing hormone receptor | GnRH_1_ | MANNASLEQDPNHCSAINNSIPLIQGKLPTLTVSGKIR |
| GPER1 | G-protein coupled estrogen receptor 1 | GPER | MDVTSQARGVGLEMYPGTAQPAAPNTTSPELNLSHPLLGTALANGTGELSEHQQYVIGLFLS |
| GPR12 | G-protein coupled receptor 12 | GPR12 | MNEDLKVNLSGLPRDYLDAAAAENISAAVSSRVPAVEPEPELVVNPW |
| GPR182 | G-protein coupled receptor 182 | GPR182 | MSVIPSSRPVSTLAPDNDFREIHNWTELLHLFNQTFSDCHMELNENTKQVVLF |
| GPR20 | G-protein coupled receptor 20 | GPR20 | MPSALSMRPWDAALPNTTAAAWTNGSVPEMPLFHHFARLDEELQAT |
| GRPR | Gastrin-releasing peptide receptor | BB_2_ | MAPNNCSHLNLDVDPFLSCNDTFNQSLSPPKMDNWFHPG |
| HCRTR1 | Orexin receptor type 1 | OX_1_ | MEPSATPGPQMGVPTGVGDPSLVPPDYEEEFLSYLWRDYLYPKQYE |
| HRH3 | Histamine H3 receptor | H_3_ | MERAPPDGPLNASGALAGEAAAAGGARGFSAAWTAVLAA |
| HTR1A | 5-hydroxytryptamine receptor 1A | 5-HT_1A_ | MDVLSPGQGNNTTSPPAPFETGGNTTGISDVTVSYQ |
| HTR1B | 5-hydroxytryptamine receptor 1B | 5-HT_1B_ | MEEPGAQCAPPPPAGSETWVPQANLSSAPSQNCSAKD |
| HTR1D | 5-hydroxytryptamine receptor 1D | 5-HT_1D_ | MSPLNQSAEGLPQEASNRSLNATETSEAWDPRTLQAL |
| HTR2A | 5-hydroxytryptamine receptor 2A | 5-HT_2A_ | MDILCEENTSLSSTTNSLMQLNDDTRLYSNDFNSGEANTSDAFNWTVDSENRTNLSCEGCLSPSCLSLL |
| HTR2B | 5-hydroxytryptamine receptor 2B | 5-HT_2B_ | MALSYRVSELQSTIPEHILQSTFVHVISSNWSGLQTESIPEEMKQIV |
| KISS1R | KiSS-1 receptor | kisspeptin | MAAEATLGPNVSWWAPSNASGCPGCGVNASDGPGSAPRPLDAWLVP |
| MAS1L | Mas-related G-protein coupled receptor | MAS1L | MVWGKICWFSQRAGWTVFAESQISLSCSLCLHSGDQEAQNPNLVSQLCGVFLQNETN |
| MC3R | Melanocortin receptor 3 | MC_3_ | MNSSCCLSSVSPMLPNLSEHPAAPPASNRSGSGFCEQ |
| MLNR | Motilin receptor | motilin | MGSPWNGSDGPEGAREPPWPALPPCDERRCSPFPL |
| MRGPRD | Mas-related G-protein coupled receptor member D | MRGPRD | MNQTLNSSGTVESALNYSRGSTVHTAYLVLSSL |
| MRGPRX2 | Mas-related G-protein coupled receptor member X2 | MRGPRX2 | MDPTTPAWGTESTTVNGNDQALLLLCGKETLIP |
| MTNR1B | Melatonin receptor type 1B | MT_2_ | MSENGSFANCCEAGGWAVRPGWSGAGSARPSRTPRPP |
| NPBWR1 | Neuropeptides B/W receptor type 1 | NPBW_1_ | MHNLSLFEPGRGNVSCGGPFLGCPNESNPAPLPLPQPLA |
| NPSR1 | Neuropeptide S receptor | NPS | MPANFTEGSFDSNGTGQMLDSSPVACTETVTFTEVVEGKEWGSFYYSFKTEQ |
| NPY1R | Neuropeptide Y receptor type 1 | Y_1_ | MNSTLFSKVENHSIHYNASENSPLLAFENDDCH |
| NPY2R | Neuropeptide Y receptor type 2 | Y_2_ | MGPIGAEADENQTVEEMKVEQYGPQTTPRGELVPDPEPELIDSTKLIEVQV |
| OPN5 | Opsin-5 | OPN5 | MALNHTALPQDERLPHYLRDGDPFASKLSWEAD |
| OPRL1 | Nociceptin receptor | NOP | MEPLFPAPFWEVIYGSHLQGNLSLLSPNHSLLPPHLLLNASHGAFLPL |
| OXGR1 | 2-oxoglutarate receptor 1 | oxoglutarate | MIETLDSPANDSDFLDYITALENCTDEQISFKMQYLP |
| P2RY13 | P2Y purinoceptor 13 | P2Y_13_ | MTAAIRRQRELSILPKVTLEAMNTTVMQGFNRSERCPRDTRIVQLVFPA |
| PRLHR | Prolactin-releasing peptide receptor | PrRP | MASLPTQGPAAPDFFNGLLPASSSPVNQSSETVVGNGSAAGPGSQAITPFQSLQLVHQLKGL |
| PROKR1 | Prokineticin receptor 1 | PKR_1_ | METTMGFMDDNATNTSTSFLSVLNPHGAHATSFPFNFSYSDYDMPLDEDEDVTNSRTFFAAK |
| PROKR2 | Prokineticin receptor 2 | PKR_2_ | MAAQNGNASFPANFSIPQEHASSLPFNFSYDDYDLPLDEDEDMTKTQTFFAAK |
| PTGDR2 | Prostaglandin D2 receptor 2 | DP_2_ | MANITLKPLCPLLEEMVQLPNHSNSSLRYIDHVS |
| RHO | Rhodopsin | rhodopsin | MNGTEGPNFYVPFSNKTGVVRSPFEYPQYYLAE |
| S1PR1 | Sphingosine 1-phosphate receptor 1 | S1P_1_ | MGPTSVPLVKAHRSSVSDYVNYDIIVRHYNYTGKLNISADKENSIK |
| SSTR3 | Somatostatin receptor type 3 | SST_3_ | MAAVTYPSSVPTTLDPGNASSAWPLDTSLGNASAGTSLAGLAVSG |
| SSTR5 | Somatostatin receptor type 5 | SST_5_ | MEPLSLTSTPSWNASAASSSSHNWSLVDPVSPMGA |

**Supplementary Table 3. Amino acid sequences of the HiBiT-tagged TAS2R constructs used in the cell surface expression study**

Signal sequence (where X denotes amino acid sequence of the signal sequence in Supplementary Table 2)

Linker

HiBiT tag

| **Bitter taste receptor** | **Amino acid sequence** |
| --- | --- |
| TAS2R20 | XEFGGGSGGSSSGGVSGWRLFKKISGGSGGGGSGGSSSGGVDMSFLHIVFSILVVVAFILGNFANGFIALINFIAWVKRQKISSADQIIAALAVSRVGLLWVILLHWYSTVLNPTSSNLKVIIFISNAWAVTNHFSIWLATSLSIFYLLKIVNFSRLIFHHLKRKAKSVVLVIVLGSLFFLVCHLVMKHTYINVWTEECEGNVTWKIKLRNAMHLSNLTVAMLANLIPFTLTLISFLLLIYSLCKHLKKMQLHGKGSQDPSTKIHIKALQTVTSFLILLAIYFLCLIISFWNFKMRPKEIVLMLCQAFGIIYPSFHSFILIWGNKTLKQTFLSVLWQVTCWAKGQNQSTP |
| TAS2R38 (PAV) | XEFGGGSGGSSSGGVSGWRLFKKISGGSGGGGSGGSSSGGVDLTLTRIRTVSYEVRSTFLFISVLEFAVGFLTNAFVFLVNFWDVVKRQPLSNSDCVLLCLSISRLFLHGLLFLSAIQLTHFQKLSEPLNHSYQAIIMLWMIANQANLWLAACLSLLYCSKLIRFSHTFLICLASWVSRKISQMLLGIILCSCICTVLCVWCFFSRPHFTVTTVLFMNNNTRLNWQNKDLNLFYSFLFCYLWSVPPFLLFLVSSGMLTVSLGRHMRTMKVYTRNSRDPSLEAHIKALKSLVSFFCFFVISSCAAFISVPLLILWRDKIGVMVCVGIMAACPSGHAAVLISGNAKLRRAVMTILLWAQSSLKVRADHKADSRTLC |
| TAS2R50 | XEFGGGSGGSSSGGVSGWRLFKKISGGSGGGGSGGSSSGGVDITFLYIFFSILIMVLFVLGNFANGFIALVNFIDWVKRKKISSADQILTALAVSRIGLLWALLLNWYLTVLNPAFYSVELRITSYNAWVVTNHFSMWLAANLSIFYLLKIANFSNLLFLHLKRRVRSVILVILLGTLIFLVCHLLVANMDESMWAEEYEGNMTGKMKLRNTVHLSYLTVTTLWSFIPFTLSLISFLMLICSLCKHLKKMQLHGEGSQDLSTKVHIKALQTLISFLLLCAIFFLFLIVSVWSPRRLRNDPVVMVSKAVGNIYLAFDSFILIWRTKKLKHTFLLILCQIRC |

**Supplementary Table 4.** Projected EC_50_ of compounds that produced a partial dose-response curve when tested in transfected cells expressing both Gα16-gust44 and mt-clytin II.

| **Compound** | **EC_50_ value** |
| --- | --- |
| Brucine | 2.3 M |
| Chloroquine diphosphate | > 5 M |
| Calcium chloride | 34 mM |
| Zinc sulphate | 2.8 M |
| Sinigrin | >5 M |

**Supplementary Table 5.** Summary of the predicted length of extracellular N-terminus and *N*-glycosylation sites in TAS2Rs.

| **Uniprot ID** | **Bitter taste receptor** | **Predicted length of extracellular N-terminus (number of amino acid residues)** | **N-glycosylation sites (amino acid position)*** |
| --- | --- | --- | --- |
| Q9NYW7 | TAS2R1 | 9 | 163 |
| Q9NYW6 | TAS2R3 | 6 | 166 |
| Q9NYW5 | TAS2R4 | 9 | 164, 165, 169 |
| Q9NYW4 | TAS2R5 | 1 | 155 |
| Q9NYW3 | TAS2R7 | 9 | 167, 175 |
| Q9NYW2 | TAS2R8 | 7 | 167 |
| Q9NYW1 | TAS2R9 | 9 | 164 |
| Q9NYW0 | TAS2R10 | 6 | 92, 158 |
| Q9NYV9 | TAS2R13 | 7 | 162, 166 |
| Q9NYV8 | TAS2R14 | 7 | 153, **162**, **171** |
| Q9NYV7 | TAS2R16 | 1 | 80, **163** |
| P59542 | TAS2R19 | 1 | 161 |
| P59543 | TAS2R20 | 6 | 161, 176 |
| P59541 | TAS2R30 | 1 | 161, 176 |
| P59538 | TAS2R31 | 2 | 161 |
| P59533 | TAS2R38 | 17 | 89, 178 |
| P59534 | TAS2R39 | 30 | 185, 194 |
| P59535 | TAS2R40 | 14 | 170, 179 |
| P59536 | TAS2R41 | 7 | 167 |
| Q7RTR8 | TAS2R42 | 7 | 163 |
| P59537 | TAS2R43 | 1 | 161, 176 |
| P59539 | TAS2R45 | 1 | 161 |
| P59540 | TAS2R46 | 1 | **161**, 176 |
| P59544 | TAS2R50 | 1 | 161 |
| P59551 | TAS2R60 | 7 | 179 |

*Numbers in bold indicate Asn residues that were reported to be *N*-glycosylated.^1^

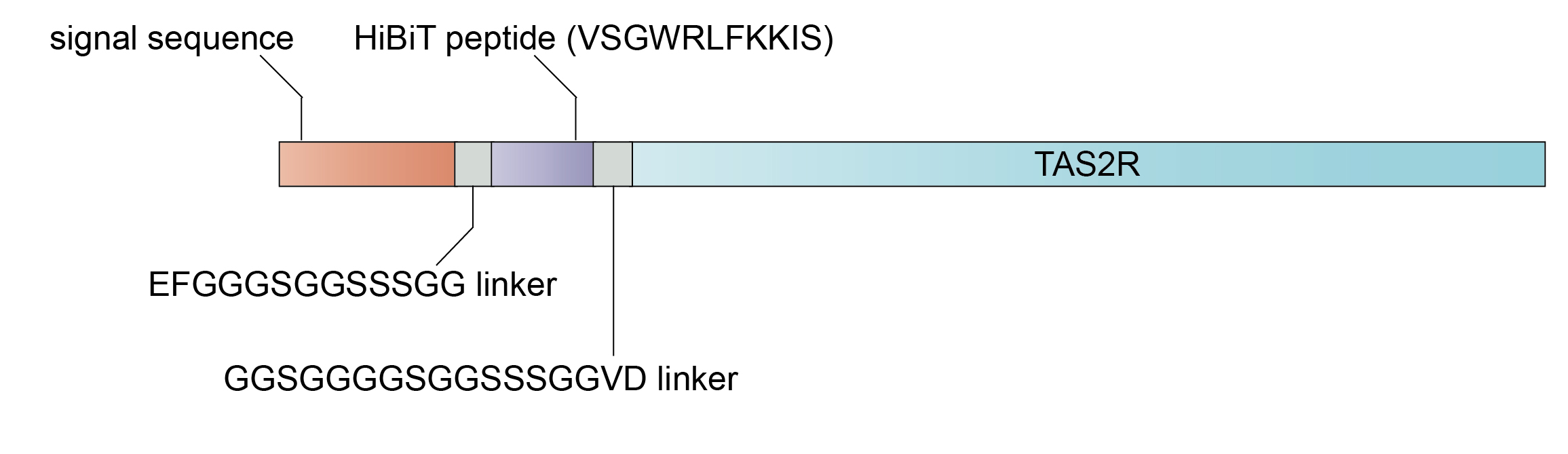

**Supplementary Figure 1.** The TAS2R construct used in the cell surface expression HiBiT assay consisted of an N-terminal signal sequence fused to a HiBiT peptide, which is flanked by EFGGGSGGSSSGG and GGSGGGGSGGSSSGGVD linkers.

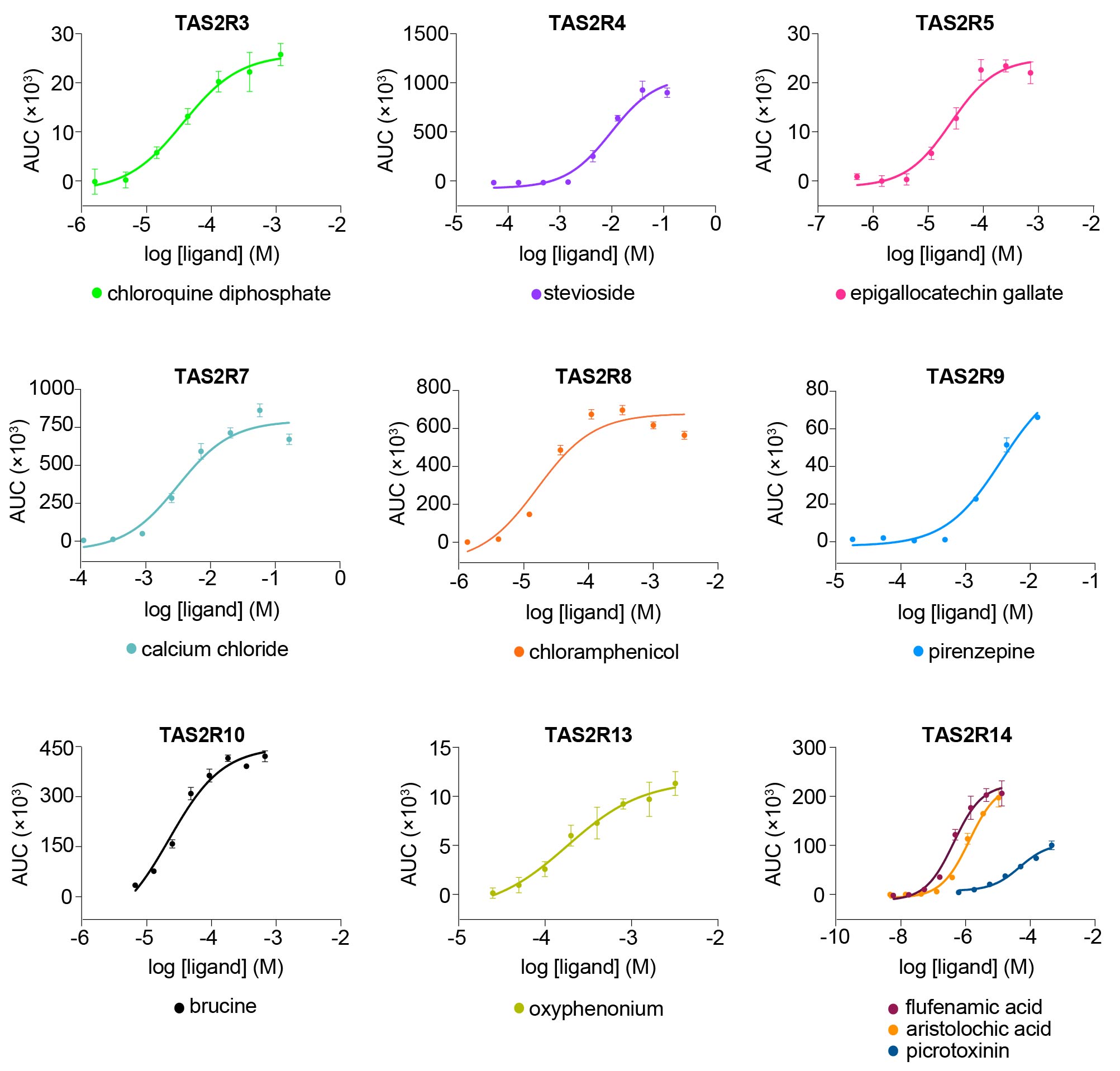

**Supplementary Figure 2. (continued)**

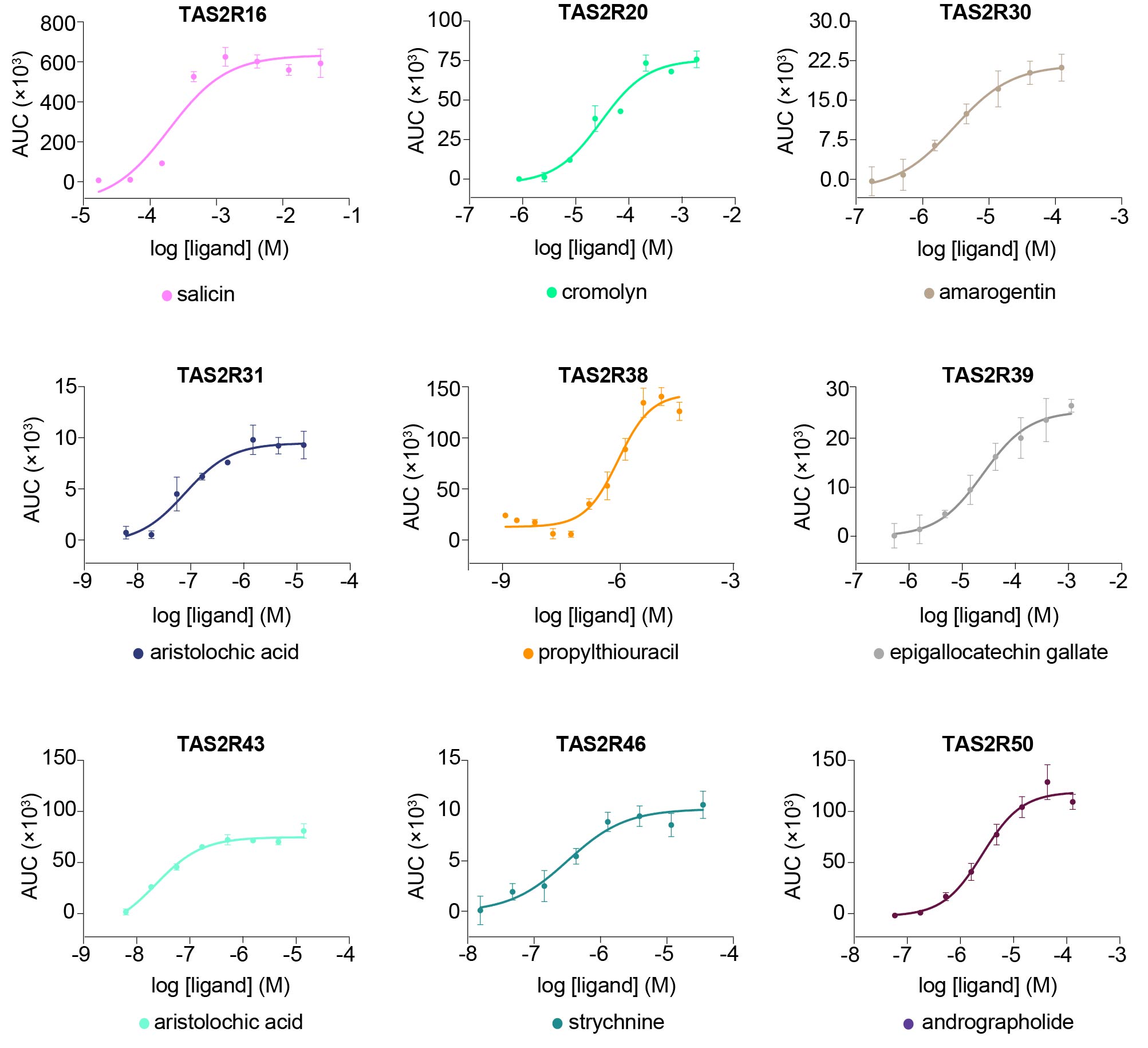

**Supplementary Figure 2.** Concentration-response curves of TAS2Rs upon stimulation with their cognate agonists in the bioluminescence-based intracellular calcium release assay in 293AD cells. Data points are shown as mean ± s.e.m. from a representative experiment out of three independent biological replicates performed in technical quadruplicates.

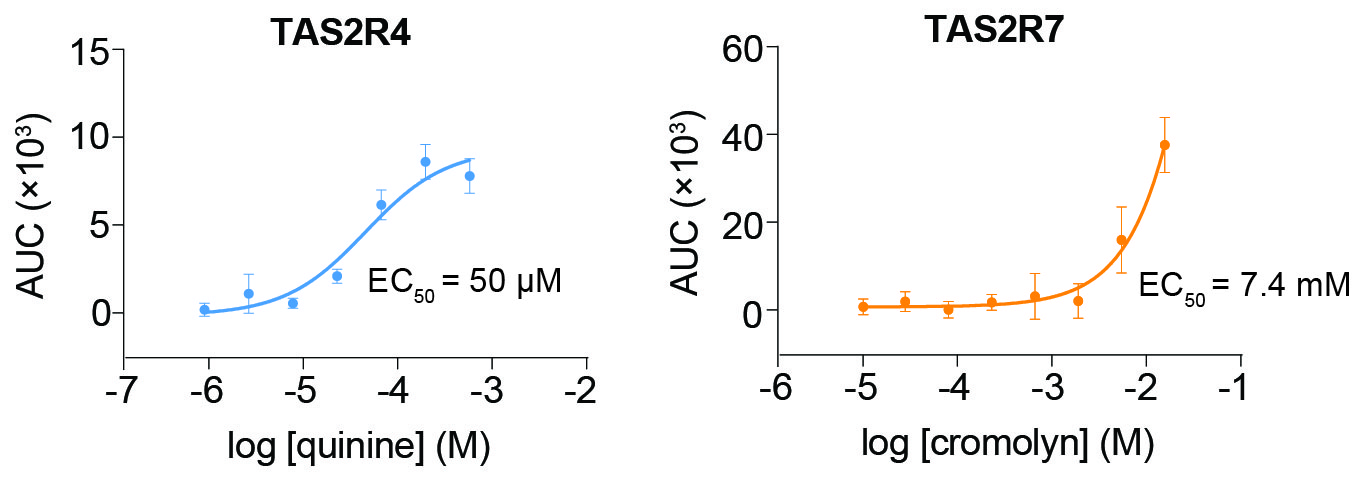

**Supplementary Figure 3**. Concentration-response curves of TAS2R4/7 upon stimulation with their agonists in the bioluminescence-based intracellular calcium release assay in AD-293 cells. The potency value of cromolyn (EC_50_ = 7.4 mM) obtained was similar to that reported in literature (EC_50_ = 5.9-6.67 mM).^2^ While no potency value was reported for quinine against TAS2R4, our experimentally derived potency value (EC_50_ = 50 μM) was close to its reported minimal effective concentration (10 μM) that elicited response from TAS2R4-expressing cells.^3^ Data points are shown as mean ± s.e.m. from a representative experiment out of two independent biological replicates performed in technical quadruplicates.

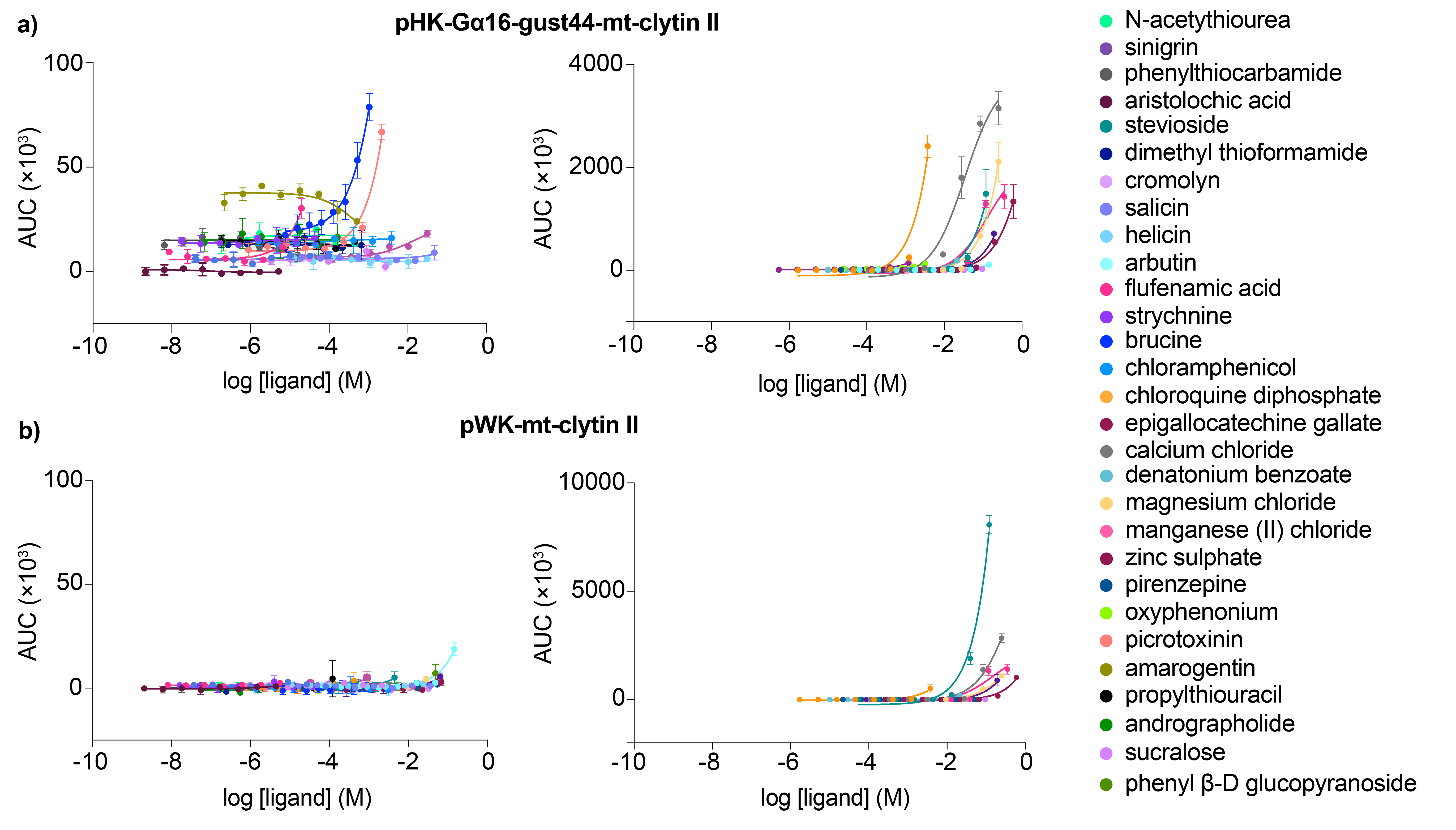

**Supplementary Figure 4**. Concentration-response curves of TAS2R agonists in cells transfected to express either a) Gα16-gust44 and mt-clytin II, or b) solely mt-clytin II. Data points are shown as mean ± s.e.m. from a representative experiment out of three independent biological replicates performed in technical quadruplicates.

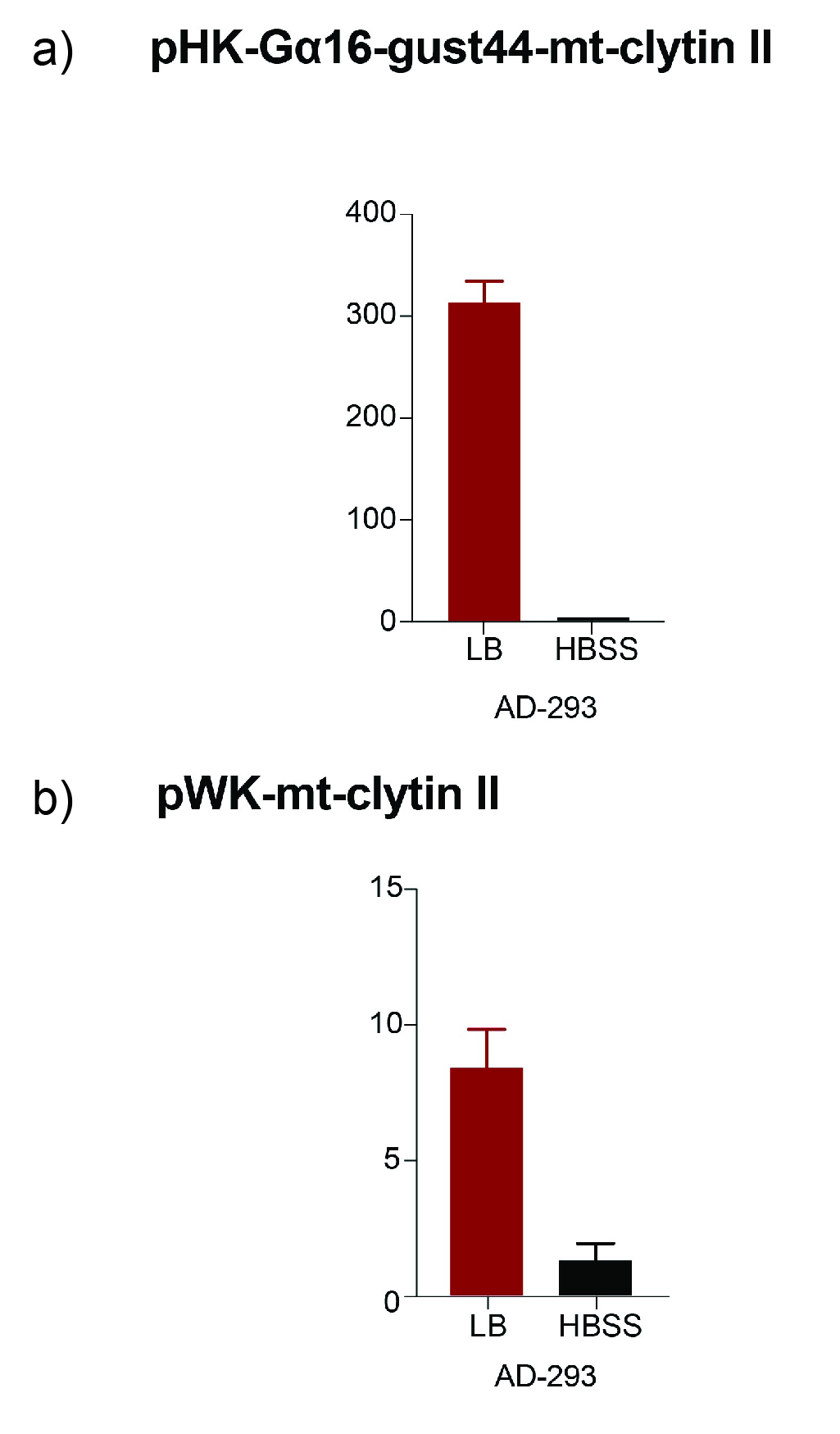

**Supplementary Figure 5**. Activation of calcium responses by LB medium and HBSS in AD-293 cells transfected to express a) both Gα16-gust44 and mt-clytin II, or b) solely mt-clytin II. Data points are shown as mean ± s.e.m. from a representative experiment out of two independent biological replicates performed in technical quadruplicates.
